# Exposure to high-sugar diet induces transgenerational changes in sweet sensitivity and feeding behavior via H3K27me3 reprogramming

**DOI:** 10.1101/2023.01.15.524137

**Authors:** Jie Yang, Ruijun Tang, Shiye Chen, Yinan Chen, Liudeng Zhang, Kai Yuan, Rui Huang, Liming Wang

**Affiliations:** Life Sciences Institute, Zhejiang University, Hangzhou, 310058, Zhejiang, China.; Hunan Key Laboratory of Molecular Precision Medicine, Department of Neurosurgery, Xiangya Hospital, and Hunan Key Laboratory of Medical Genetics, School of Life Sciences, Central South University, Changsha, Hunan 410008, China; The Biobank of Xiangya Hospital, Xiangya Hospital, Central South University, Changsha, Hunan 410008, China; Center for Neurointelligence, School of Medicine, Chongqing University, Chongqing, 400030, China; Institute of Molecular Physiology, Shenzhen Bay Laboratory, Shenzhen, 518055, China.

## Abstract

Human health is facing a host of new threats linked to unbalanced diets, including high sugar diet (HSD), which contributes to the development of both metabolic and behavioral disorders. Studies have shown that diet-induced metabolic dysfunctions can transmit to multiple generations of offspring and exert long-lasting health burden. Meanwhile, whether and how diet-induced behavioral abnormalities can be transmitted to the offspring remain largely unclear. Here, we showed that ancestral HSD exposure suppressed sweet sensitivity and feeding behavior in the offspring in *Drosophila*. These behavioral deficits were transmitted through the maternal germline and companied by the enhancement of H3K27me3 modifications. PCL-PRC2 complex, a major driver of H3K27 trimethylation, was upregulated by ancestral HSD exposure, and disrupting its activity eliminated the transgenerational inheritance of sweet sensitivity and feeding behavior deficits. Elevated H3K27me3 inhibited the expression of a transcriptional factor Cad and suppressed sweet sensitivity of the sweet-sensing gustatory neurons, reshaping the sweet perception and feeding behavior of the offspring. Taken together, we uncovered a novel molecular mechanism underlying behavioral abnormalities across multiple generations of offspring upon ancestral HSD exposure, which would contribute to the further understanding of long-term health risk of unbalanced diet.

## INTRODUCTION

Dietary factors play a critical role in regulating multiple biological processes and influencing animal metabolism and behavior. For example, dietary restriction extends lifespan through metabolic regulation (Anson et al., 2003; Wu et al., 2019), while high-fat diet (HFD) and high-sugar diet (HSD) lead to obesity and various metabolic dysfunctions (Birse et al., 2010; Buettner et al., 2007; Palanker Musselman et al., 2011). Evidence has also emerged indicating that dietary factors impact gene expression through epigenetic modifications, which may contribute to these metabolic syndromes (Park et al., 2017). In addition to the direct effects of dietary changes within the same generation of animals, dietary changes may also lead to alterations in the germline cells which exerts long-lasting effects in the following generations. Studies on individuals who were born during the Dutch and Chinese famine demonstrate that prenatal exposure to undernutrition environments causes overweight and insulin resistance in the offspring (Heijmans et al., 2008; Li & Lumey, 2017; Ravelli et al., 1998; Tobi et al., 2014). Animal models such as worms, flies, and mice also indicate that exposure to abnormal diets induces various transgenerational metabolic disorders, including diabetes, obesity, hyperlipidemia, and so forth (Dunn & Bale, 2011; Huypens et al., 2016; Somer & Thummel, 2014; Stegemann & Buchner, 2015; Wei et al., 2014).

Such transgenerational inheritance upon dietary changes is thought to be mediated by several epigenetic factors, including DNA methylation, ncRNA, and histone modifications (Bohacek & Mansuy, 2015; Heard & Martienssen, 2014; Miska & Ferguson-Smith, 2016; Skvortsova et al., 2018). For example, studies in *C. elegans* demonstrate that starvation alters organismal metabolism spanning three subsequent generations via small RNAs (Rechavi et al., 2014), and HFD induces lipid accumulation signals transmitting to multiple generations through H3K4me3 modifications (Wan et al., 2022). Similarly, previous reports in HFD mouse model show that *in utero* exposure to HFD causes a metabolic syndrome through epigenetic modifications of adiponectin and leptin signaling, and that sperm tsRNA signaling contributes to intergenerational inheritance of an acquired metabolic disorder (Chen et al., 2016; Masuyama & Hiramatsu, 2012). Dietary factors can also alter animal behaviors. For example, HFD affects the feeding and cognitive behaviors of mice (Arnold et al., 2014; Pendergast et al., 2013). In human studies, children of famine survivors had higher chances to develop psychological trauma or insanity (Kelly, 2019; Li et al., 2015; Painter et al., 2006), which implies that an abnormal diet may lead to behavioral disorders with transgenerational inheritance.

The fruit flies *Drosophila melanogaster* is a valuable model for studying the transgenerational inheritance of animal behaviors. The effects of diet changes on various fly behaviors have been demonstrated in flies. For example, starvation increases files’ locomotion and food-seeking behavior (Yu et al., 2016), and HSD reshapes sweet perception and promotes feeding (May et al., 2019). Moreover, there is accumulating evidence of diet-induced transgenerational inheritance in *Drosophila*. For example, changes in dietary yeast concentrations induce transgenerational somatic rDNA instability and copy number reduction (Aldrich & Maggert, 2015). HFD exposure induces transgenerational cardiac lipotoxicity through H3K27me3 modifications (Guida et al., 2019). A low protein diet leads to transgenerational reprogramming of lifespan through E(z)-mediated H3K27me3 modifications (Xia et al., 2016). There is also evidence that behavioral changes in *Drosophila* can be transmitted to subsequent generations: exposure to predatory wasps leads to transgenerational ethanol preference via maternal NPF repression (Bozler et al., 2019).

HSD results in many physiological responses and metabolic/behavioral disorders in the same generation of *Drosophila* (Chen et al., 2021; Chng et al., 2017; May et al., 2019; van Dam et al., 2020). Some metabolic changes, such as obese- and diabetes-like phenotypes, can be passed on to their offspring through germline epigenetic alternations (Buescher et al., 2013; Karunakar et al., 2019; Öst et al., 2014; Palanker Musselman et al., 2011; van Dam et al., 2020). However, the possibility and mechanisms of transgenerational inheritance of behavioral changes upon HSD exposure are far less studied in fruit flies.

In this study, we found that HSD induced metabolic and behavioral dysfunctions as previously reported, and discovered that HSD suppressed sweet sensitivity and feeding behavior in the offspring. Furthermore, ChIP-seq data revealed that this transgenerational behavioral change was mediated by up-regulated H3K27me3 modifications transmitted through the maternal germline. More specifically, we identified that ancestral HSD exposure elevated H3K27me3 levels in the promoter region of *cad* gene, resulting in a reduction in its mRNA expression in the sweet-sensing gustatory neurons of offspring, eventually reshaping the sweet perception and feeding behavior. Taken together, our study uncovered a novel molecular mechanism underlying the transgenerational behavioral changes upon ancestral HSD exposure, and shed light on the understanding of long-term health risks of dietary abnormalities in human.

## RESULTS

### HSD feeding suppressed sweet sensitivity and feeding behavior spanning multiple generations

Previous works have shown that ancestral exposure of abnormal diets (such as HFD and HSD) leads to various metabolic dysfunctions in the offspring, including cardiac lipotoxicity, diabetes, and obesity (Chen et al., 2022; Guida et al., 2019; Kaspar et al., 2020; Wan et al., 2022). However, whether ancestral experience exerted transgenerational behavioral changes was still unclear. To address this question, we used *Drosophila melanogaster* as a model system to examine the potential transgenerational behavioral effect of HSD.

Wild-type flies were fed with HSD from the embryo stage to adulthood (termed HSD-F0 flies). Fresh embryos of HSD-F0 flies were transferred back to normal diet (ND) and raised on ND to adulthood (termed HSD-F1 flies). These HSD-F1 flies were further raised and mated on ND to generate multiple generations of offspring (HSD-F2 to F5) (*Figure 1A*). Flies continuously raised on ND without any HSD exposure were used as ND controls. Notably, HSD-F1 to F5 flies and ND-fed control flies were all raised on ND food from the embryo stage, thus any metabolic and behavioral differences among them were likely attributed to ancestral exposure to HSD and its transgenerational effect on offspring.

**Figure 1.**
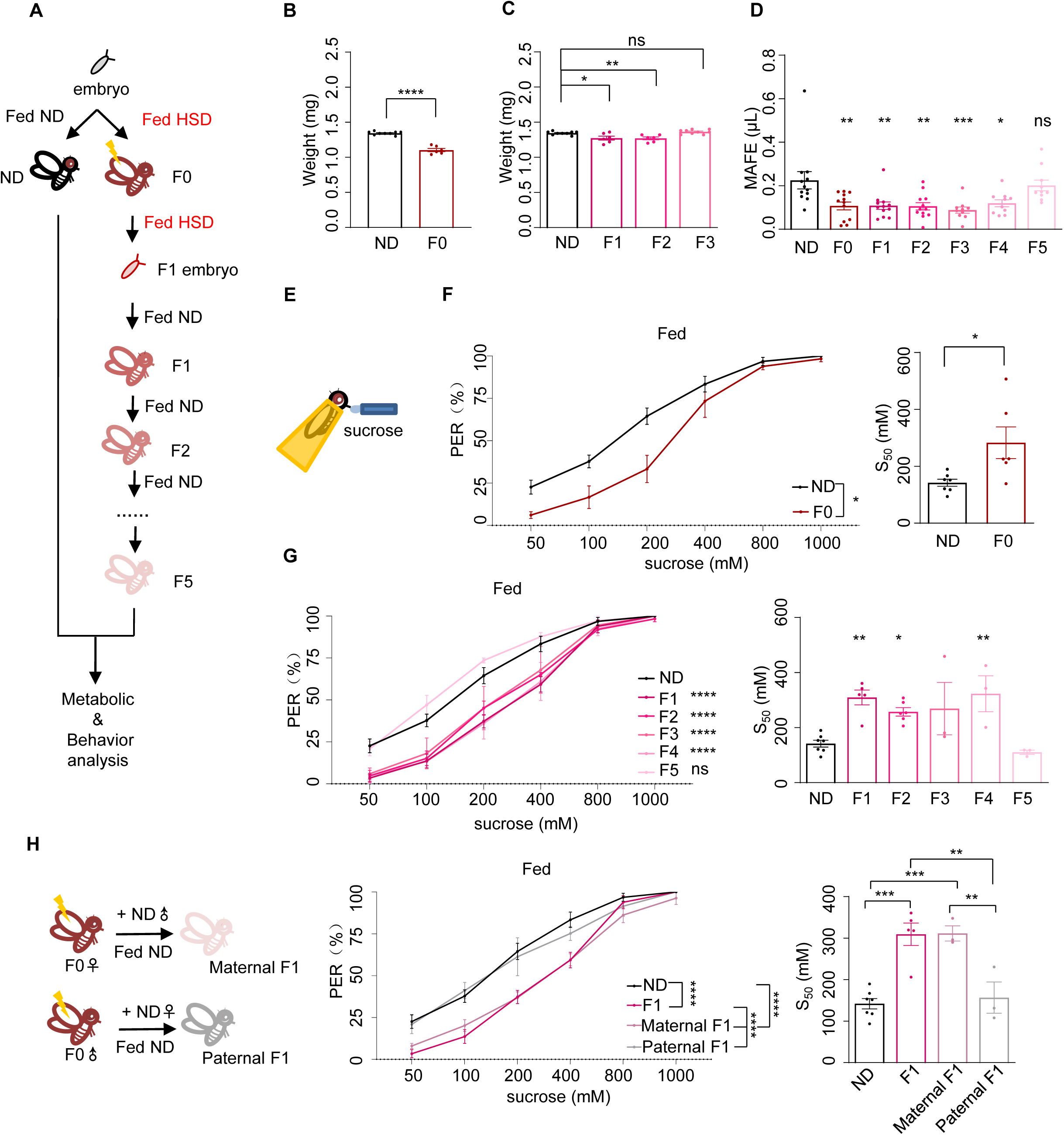
Ancestral HSD exposure decreased sweet sensitivity and feeding behavior across multiple generations of offspring. A. The illustration of experimental design for Figure 1B-H. The embryos of wild-type *Canton-S* flies were collected and fed with ND (black, referred to ND controls) or HSD (red, referred to HSD-F0) until maturity. HSD-F0 flies were mated to produce the next generation (HSD-F1). The embryos of HSD-F1 flies were transferred to ND right after egg laying and kept on ND to adulthood. HSD-F1 flies were mated to propagate multiple generations of offspring (HSD-F2 to F5) on ND diet. These flies were prepared for different metabolic and behavioral assays. B-C. The body weight of individual flies from different treatment groups (n = 6 biological replicates, each containing 5 flies). D. Volume of 400 mM sucrose consumed by individual flies using the MAFE assay (n = 12). E. Schematic illustration of the PER assay. F-H. Fractions of flies showing PER responses to different concentrations of sucrose (n = 3-6 biological replicates, each containing 8-12 flies). The S_50_ indicated the sucrose concentration that induced PER responses in 50% of the tested flies. Data were shown as means ± SEM. ns P>0.05; * P<0.05; ** P < 0.01; *** P < 0.001; **** P < 0.0001.

To investigate the consequences of ancestral HSD exposure, we first assayed multiple physiological and metabolic parameters of HSD-F0 and HSD-F1 flies versus ND-fed controls. Compared to ND flies, both HSD-F0 and HSD-F1 flies exhibited decreased body weight (*Figure 1B-C*). In contrast, their glycogen and triglyceride storage levels, as well as trehalose levels, the major circulating sugar in the fly hemolymph, were elevated (*Figure 1-figure supplement 1A-C*). Hyperglycemia together with weight loss was a sign of insulin signaling dysfunction. We therefore quantified the mRNA levels of two important insulin-like molecules in flies, *Drosophila* Insulin-Like Peptide 2 (DILP2) and DILP5, both of which were released by insulin-producing cells in the fly brain upon nutrient uptake (Ikeya et al., 2002). As expected, the expression levels of these genes were down-regulated in HSD-F0 and HSD-F1 flies (*Figure 1-figure supplement 1D-E*). These data suggest that HSD feeding leads to the development of a diabetes-like phenotype in the same generation (HSD-F0) and that such phenotype can be passed on to their offspring (HSD-F1), as previously reported (Buescher et al., 2013; Palanker Musselman et al., 2011; van Dam et al., 2020).

Next, we asked whether ancestral HSD exposure resulted in behavioral abnormalities and whether this effect could also be transmitted to the offspring. Given that insulin signaling played an important role in feeding regulation (Porte et al., 2002), we first measured total food consumption by the Capillary Feeder (CAFE) assay in HSD-F0 flies (Ja et al., 2007). As previously reported, HSD-F0 flies exhibited significantly increased food consumption compared to ND-fed controls in a 24-hour duration; similar increases were observed in HSD-F1 and HSD-F2 flies, too (*Figure 1-figure supplement 2A*). However, when we used our previously developed Manual Feeding (MAFE) assay (Qi et al., 2015) to measure the volume of ingested food by individual flies during the course of a single meal, we found that HSD-F0 flies exhibited significantly decreased food consumption, and that the suppression of meal size was transmitted through multiple generations (*Figure 1D*).

The difference between these two feeding assays was that in the MAFE assay flies were immobilized and presented with micro capillaries filled with liquid food, whereas in the CAFE assay flies were free-moving and could decide when to feed. A major determinant of the readout of the MAFE assay was whether flies were responsive to the presented food and were willing to expend their proboscis to initiate a meal, and that of the CAFE assay was flies’ overall energy need. Thus, the discrepancy between the results from the CAFE assay and the MAFE assay suggests that upon HSD exposure, flies’ overall energy expenditure is elevated while their sweet perception is inhibited, hence their increased food consumption but reduced meal size.

To test this hypothesis, we used proboscis extension reflex (PER), a behavioral component of feeding initiation (Inagaki et al., 2012), to examine sweet sensitivity of these flies (*Figure 1E*). We found that both starved and fed HSD-F0 flies showed reduced PER responses to various concentrations of sucrose compared to ND controls (*Figure 1F, Figure 1-figure supplement 2B*), and such effect could transmit to following generations till HSD-F5 (*Figure 1G, Figure 1-figure supplement 2C*).

Since HSD exposure modulated both metabolism and feeding behavior in the progeny, it was possible that these two effects were connected, i.e. altered metabolism upon ancestral HSD exposure affected feeding behavior. However, there were several lines of evidence suggesting against this possibility. For example, the body weight and nutrient storage (triglyceride, glycogen, and circulating trehalose) returned to normal in HSD-F2 or HSD-F3 flies (*Figure 1C, Figure 1-figure supplement 1A-C*), whereas HSD-F2 to HSD-F4 flies still exhibited reduced meal size (*Figure 1D*) and PER responses to sucrose *(Figure 1G*). In addition, DILP2 and DILP5 expression levels were both reduced by HSD exposure from HSD-F0 to HSD-F2 flies (*Figure 1-figure supplement 1D-E*), which were known feeding suppressors (Nässel et al., 2013). Therefore, it was unlikely that altered metabolism or insulin signaling were responsible for the reduction in sweet sensitivity and feeding behavior in HSD-exposed flies.

Transgenerational inheritance could be mediated by male or female gametes as shown in previous works (Daxinger & Whitelaw, 2012; Heard & Martienssen, 2014). To distinguish between these two possibilities, we performed HSD feeding in only female or male F0 flies and crossed them with ND-fed mates. As shown in *Figure 1H*, the reduction in PER was only seen in F1 flies with HSD-fed female ancestor but not with HSD-fed male ancestor. Similar results were observed in the F2 generation, that only F2 flies with HSD-F1 female ancestor exhibited reduction in PER (*Figure 1-figure supplement 2D*). These results suggest that the effect of HSD exposure is transmitted to offspring via female gametes.

### Ancestral HSD exposure elevated genome-wide H3K27me3 levels in offspring

Next, we investigated the underlying mechanism of transgenerational behavioral inheritance after ancestral HSD exposure. We focused on epigenetic regulators since it was unlikely that HSD exposure resulted in specific genetic alterations in the germline cells of ancestral flies. Despite the importance of DNA methylation in the regulation of vertebrate transgenerational inheritance, it was reported that DNA methylation in *Drosophila* was negligible and limited to the early stages of embryonic development (Lyko et al., 2000). Multiple lines of research indicated that two types of histone methylations, H3K27me3 and H3K9me3, played important roles in transgenerational inheritance in *Drosophila* (Coleman & Struhl, 2017; Wang & Moazed, 2017). In addition, piwi-interacting RNA (piRNA), an important species of non-coding RNA (ncRNA) in transgenerational inheritance of *Drosophila*, was associated with altered H3K27me3 and H3K9me3 (Le Thomas et al., 2014; Peng et al., 2016). Therefore, we asked whether ancestral HSD exposure induced alterations in post-translational modifications of H3K27 and H3K9.

We collected mitotic cycle 11 embryos of both ND and HSD-F1 flies, and performed chromatin-immunoprecipitation followed by sequencing (ChIP-seq) using antibodies against four histone modifications (H3K27me3, H3K27ac, H3K9me2, and H3K9me3) respectively (*Figure 2A*). We performed peak calling and determined the occupancy of H3K27me3, H3K27ac, H3K9me2, H3K9me3, and H3 on the fly genome. Genomic snapshots of representative target loci (Hox cluster genes: *bxd*, *Ubx,* and *Abd-A*) confirmed H3K27me3, H3K27ac, and H3K9me3 enrichment as expected (*Figure 2B*) and all replicates were highly comparable (*Figure 2-figure supplement 2A*). Meanwhile, we also observed that H3K27me3 and H3K27ac were widely spread in the genome at this stage, while H3K9me3 was preferentially localized within gene desert regions and heterochromatic regions such as centromeres and pericentromeric (data not shown). Our analysis also confirmed that H3K9me2 was nearly undetectable in euchromatic regions during this stage (*Figure 2B*).

**Figure 2.**
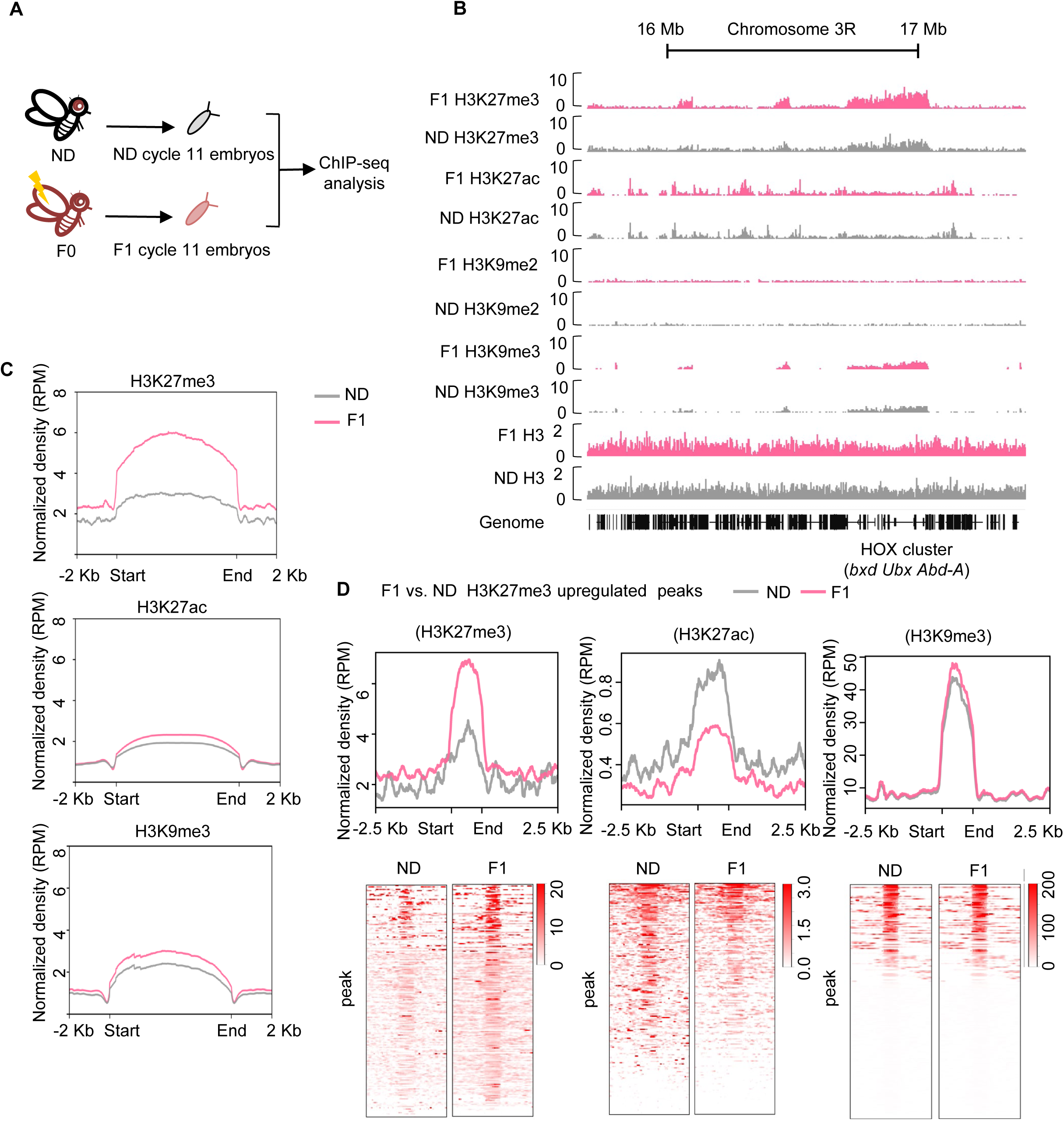
Ancestral HSD exposure increased genome-wide H3K27me3 levels in the offspring. A. The workflow of the ChIP-seq assay. ND and HSD-F1 embryos were collected from population cages at 25°C for 30 mins, and allowed to develop for 80 mins to target mitotic cycle 11. These embryos were rapidly frozen in liquid nitrogen and then used for ChIP-seq analysis. B. Genome browser view of H3K27me3, H3K27ac, and H3K9me3 density at the HOX cluster (bxd, Ubx, and Abd-A) gene regions in embryos of ND and HSD-F1 embryos. C. Average density plots showing the signal profiles of H3K27me3, H3K27ac, and H3K9me3 at their peaks plotted across a 4 kb window (± 2Kb around the start/end of signals). D. Average density plots (top) and Heatmap (bottom) showing the distribution for the changes of H3K27me3, H3K27ac and H3K9me3 signals for regions with up-regulated H3K27me3 peaks in HSD-F1 embryos, respectively. Color bar showed the Z-score value in Heatmap.

We then identified genomic regions that were significantly enriched for H3K27me3, H3K27ac, and H3K9me3 in HSD-F1 embryos using Model-based Analysis of ChIP-seq 2 (MACS2) and generated the average signal density spanning a 2 kb region at both ends. We found that upon ancestral HSD exposure, the average peak intensity of H3K27me3 modifications increased significantly (*Figure 2C*, *upper*) whereas those intensity of H3K27ac and H3K9me3 only exhibited modest increase (*Figure 2C*, *middle* and *lower*). In addition, previous work indicated that *Drosophila* oocytes transmitted repressive H3K27me3 marks to their offspring and exerted developmental impact (Zenk et al., 2017). Thus, we focused on the upregulation of H3K27me3 signals upon ancestral HSD exposure for further analysis.

To identify specific up-regulated H3K27me3 peaks between HSD-F1 and ND embryos, we performed differential peaks analysis and found that ancestral HSD exposure resulted in approximately 400 regions with H3K27me3 upregulation (log_10_ likelihood ratio>3) distributed throughout the genome and 6 H3K27me3 down-regulated regions (Supplementary file 1). The signal intensity of H3K27me3 increased remarkably in those up-regulated regions (*Figure 2D*, *left*). In these regions, the signal intensity of H3K27ac was significantly decreased (*Figure 2D*, *middle*), in line with the antagonism between H3K27ac and H3K27me3 modifications (Tie et al., 2009). Meanwhile, H3K9me3 signal in these regions remained unchanged (*Figure 2D*, *right*).

Next, we asked how ancestral HSD exposure led to H3K27me3 hypermethylation. Previous studies reported that Polycomb-like protein (Pcl) interacts with Polycomb repressive complex 2 (PRC2) to constitute a specific form of PCL-PRC2 complex, which generated high levels of H3K27me3 on specific genomic regions (Nekrasov et al., 2007). Besides Pcl, PRC2 complex was comprised of several major components, including Enhancer of zeste (E(z)), Suppressor of zeste 12 ((Su(z)12), Extra sex combs (Esc), and Chromatin assembly factor-1 (Caf-1) (*Figure 2-figure supplement 1B*) (Margueron & Reinberg, 2011). By using quantitative RT-PCR, we found that the mRNA expression of Pcl was up-regulated in HSD-F1 and HSD-F2 flies compared to ND controls, while mRNA expression of E(z), Su(z)12, Esc, and Caf-1 only exhibited modest yet insignificant changes (Figure 2-figure supplement 1C, D).

Taken together, our data suggest that ancestral HSD exposure exerts robust regulatory effects on histone modifications of the offspring, especially an elevation of H3K27me3 modifications. One possible mechanism is the up-regulation of Pcl, a key component in PCL-PRC2 complex responsible for H3K27me3 modifications. Given that genomic histone modifications undergo dynamic reprogramming during development, how such histone imprinting transmits to multiple generations of offspring remains unclear and is of significance for future studies (Heard & Martienssen, 2014).

### H3K27me3 was required for the transgenerational modulation of sweet sensitivity and feeding behavior upon ancestral HSD exposure

Since our data showed that HSD exposure elevated genome-wide H3K27me3 modifications in HSD-F1 embryos, we asked whether this inherited epigenetic change contributed to transgenerational behavioral deficits. To test it, we knocked down the expression of H3K27me3 catalytic enzyme in the PCL-PRC2 complex, E(z), to reduce H3K27me3 imprinting during embryogenesis. We used an embryo-specific GAL4 driver, *nosNGT-GAL4*, which is active during the blastoderm stage (an early developmental stage, within 4 hours of fertilization) of embryogenesis (Tracey et al., 2000). We observed that compared to transgenic controls, *nosNGT-GAL4>UAS-E(z) RNAi* flies exhibited similar PER responses to sucrose in ND vs. HSD-F1 flies, suggesting the loss of transgenerational behavioral deficits (*Figure 3A*). Knocking down Pcl, another component of PCL-PRC2 complex, generated similar effect as E(z) (*Figure 3B)*. Meanwhile, RNAi knockdown of histone deacetylase Rpd3 and H3K9me3 methylase Su(var)3-9 did not affect the transgenerational behavioral inheritance upon ancestral HSD exposure (*Figure 3C, D)*. These data indicate that H3K27me3 plays a crucial role in the transgenerational inheritance of sweet sensitivity and feeding behavior upon ancestral HSD exposure.

**Figure 3.**
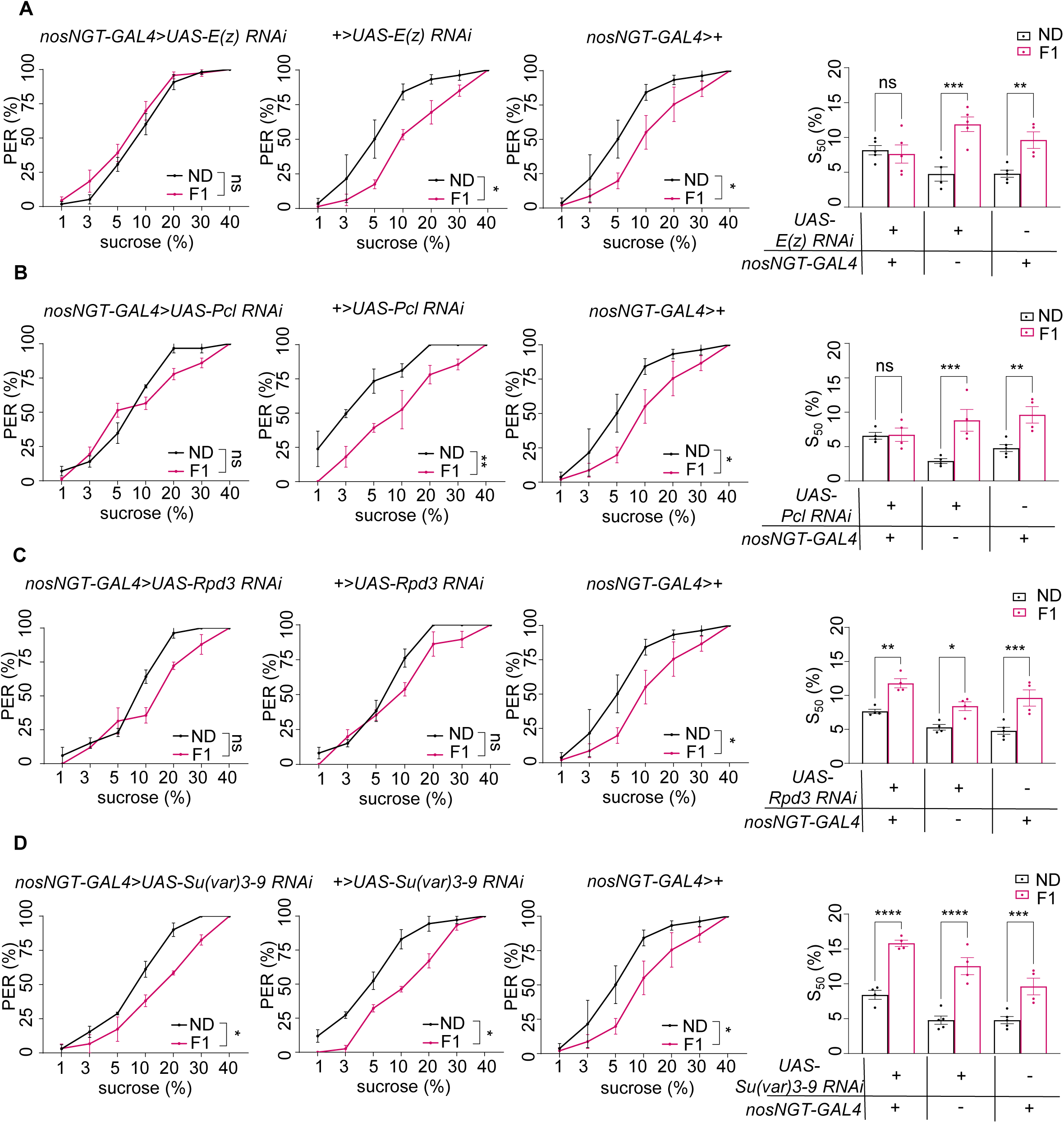
H3K27me3 modifications were essential for the transgenerational regulation of sweet sensitivity and feeding behavior upon ancestral HSD exposure. A-D. Fractions of flies of the indicated genotypes showing PER responses to sucrose (n = 3-6 biological replicates, each containing 8-12 flies). The S_50_ indicated the sucrose concentration that elicited PER responses in 50% of the tested flies. Data were shown as means ± SEM. ns P>0.05; * P<0.05; ** P < 0.01; *** P < 0.001; **** P < 0.0001.

Since our data suggested that the behavioral deficits upon ancestral HSD exposure were inherited across generations through the maternal lineage, we reasoned that H3K27me3 imprinting, the driver of the transgenerational inheritance of sweet taste deficits upon ancestral HSD exposure, was transmitted through the maternal germline. To directly test this hypothesis, we used an oocyte-specific driver GAL4, *maternal alpha-tubulin GAL4* (*Matα-tub-GAL4*), to knock down E(z) and Pcl expression during oogenesis (Hudson & Cooley, 2014). We observed that knockdown of E(z) and Pcl in female germline eliminated the suppression of PER responses in HSD-F1 flies compared to ND controls (*Figure 3-figure supplement 1*). These results confirm that the transmission of maternal H3K27me3 modifications is critical for the transgenerational inheritance of sweet sensitivity and feeding behavior.

### H3K27me3 regulated the sensitivity of sweet-sensing gustatory neurons

As we showed above, elevated H3K27me3 modifications upon ancestral HSD exposure mediated the transgenerational behavioral inheritance. We next examined whether modulating H3K27me3 levels could directly cause changes in sweet sensitivity and feeding behavior.

We tested the effect of EED226, a potent PRC2 inhibitor that directly binds to the H3K27me3 binding pocket and suppresses H3K27me3 modifications (Loh et al., 2021). In HSD-F1 to HSD-F3 flies, EED226 feeding for five consecutive days restored their PER responses to sucrose to the levels of ND controls (*Figure 4A*, *left* and *middle*). Chaetocin, a specific inhibitor of H3K9 methyltransferase Su(var)3-9 (He et al., 2012), did not rescue the sweet taste defects in HSD-F1 to HSD-F3 flies (*Figure 4A*, *right*). We further examined the effect of pharmacological inhibition of H3K27me3 by feeding HSD-F1 flies with either EED226 or A395, another histone methyltransferase inhibitor occupying the H3K27me3 binding sites (He et al., 2017), and found that both treatments restored sweet responses towards various concentrations of sucrose to the levels of ND-fed controls (*Figure 4B*). It was worth noting that knocking down E(z) and Pcl by RNAi in the embryogenesis stage (*Figure 3*) and pharmacological inhibition of H3K27me3 by two different chemicals in the adult stage (*Figure 4*) both exerted similar behavioral effects, confirming that elevated H3K27me3 modifications was indeed critical for reduced sweet sensitivity upon ancestral HSD exposure.

**Figure 4.**
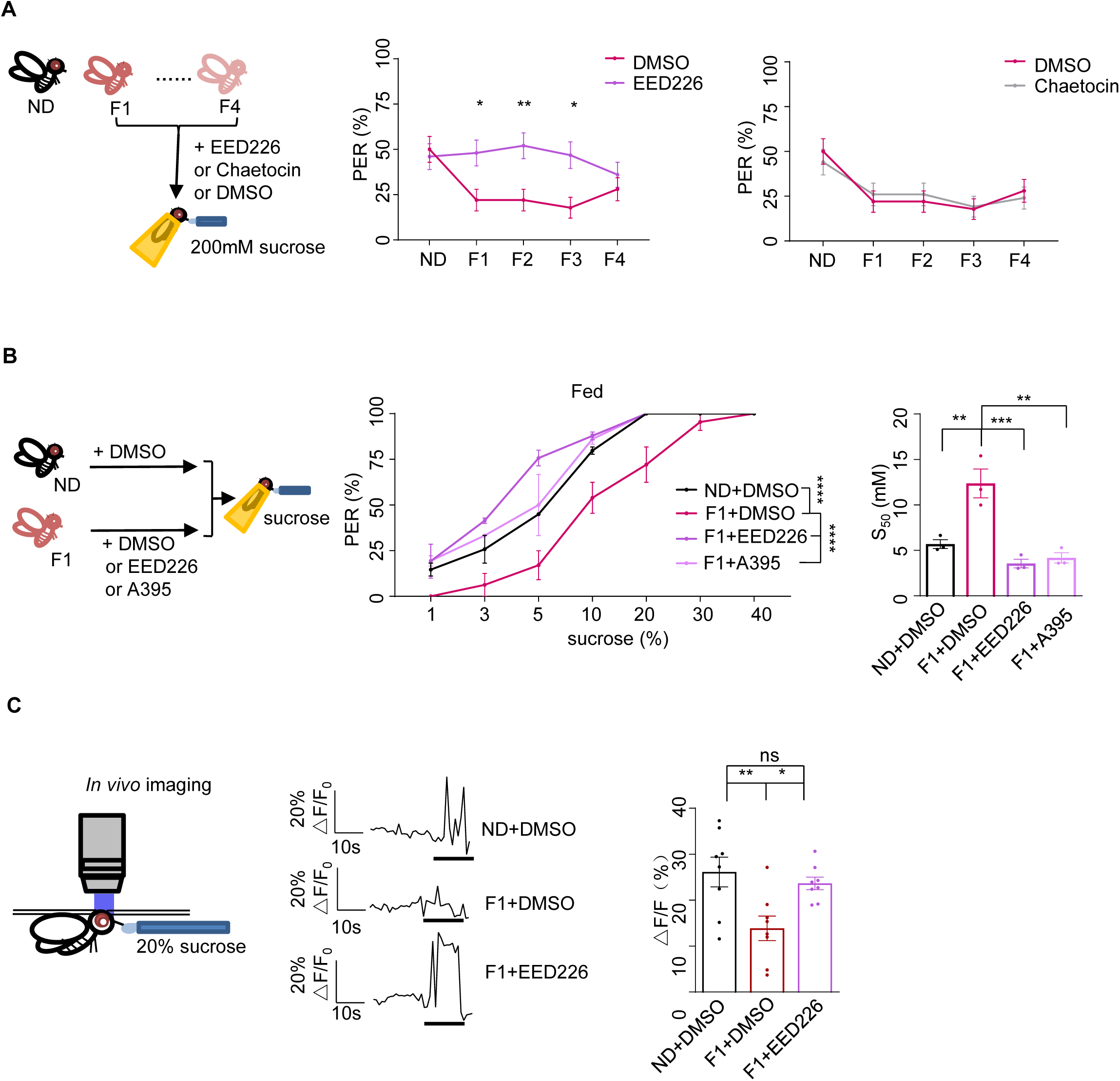
H3K27me3 modifications suppressed sweet sensitivity of Gr5a^+^ gustatory neurons. A. Fractions of flies showing PER responses to 200 mM sucrose (n = 40). Flies were fed with indicated chemicals for 5 days before the assay. B. Fractions of flies with or without EED226/A395 feeding showing PER responses to different concentrations of sucrose (n = 3-6 biological replicates, each containing 8-12 flies). The S_50_ indicated the sucrose concentration that induced PER responses in 50% of the tested flies. C. The calcium signals in Gr5a^+^ neurons in indicated flies upon 20% sucrose. Schematic diagram of *in vivo* calcium imaging was shown on *left*. Representative traces of the calcium responses were shown in *middle*. Horizontal black bars represent feeding episodes. Quantification of the calcium responses was shown on *right* (n = 7-9). Data were shown as means ± SEM. ns P>0.05; * P<0.05; ** P < 0.01; *** P < 0.001; **** P < 0.0001.

We next sought to understand the underlying neurobiological mechanism of transgenerational sweet taste deficits upon ancestral HSD exposure. Gustatory neurons expressing a gustatory receptor Gr5a played a central role in sugar perception (Dahanukar et al., 2007). We thus hypothesized that H3K27me3 modifications might modulate sweet sensitivity of Gr5a^+^ gustatory neurons. To directly test this, we ectopically expressed a genetically encoded calcium indicator GCaMP6m (Yang et al., 2018) in Gr5a^+^ neurons and conducted live calcium imaging during sucrose feeding episodes (*Figure 4C*). Indeed, HSD-F1 flies exhibited significantly reduced calcium transients upon sucrose stimulation compared to ND flies, which could be restored by EED226 feeding (*Figure 4C*).

### Cad mediated the reduction in sweet sensitivity by ancestral HSD exposure

We then asked how elevated H3K27me3 modifications altered sweet sensitivity of Gr5a^+^ neurons. Previous work reported that the formation of H3K27me3 modifications maintained the silenced state of *Drosophila* homeobox genes and transmitted such a repressive state through multiple rounds of DNA replication in the early embryos and exerted long-lasting impact into adulthood (Coleman & Struhl, 2017; Skvortsova et al., 2018). We speculated that up-regulated H3K27me3 in the early embryos could alter gene transcriptome in certain organs and affect the physiology and behavior of adults.

To include more candidate genes for further analysis, we used less stringent criteria compared to *Figure 2* (log_10_ likelihood ratio>1) and identified ∼3000 H3K27me3 hypermethylated regions in HSD-F1 flies using a window of ± 1 kb from the transcription start site (TSS) and found 340 genes corresponding to hypermethylated sites (Supplementary file 2). Since H3K27me3 often marks the downregulation of gene expression, we then asked whether and which of these genes were transcriptionally suppressed. We performed RNA-seq analysis using the head tissues of HSD-F1 vs. ND flies (*Figure 5A*) and identified a total of 136 differentially expressed genes (DEG, |log_1.5_ fold change|>1, P<0.05), including 50 down-regulated genes in HSD-F1 flies (Supplementary file 3).

**Figure 5.**
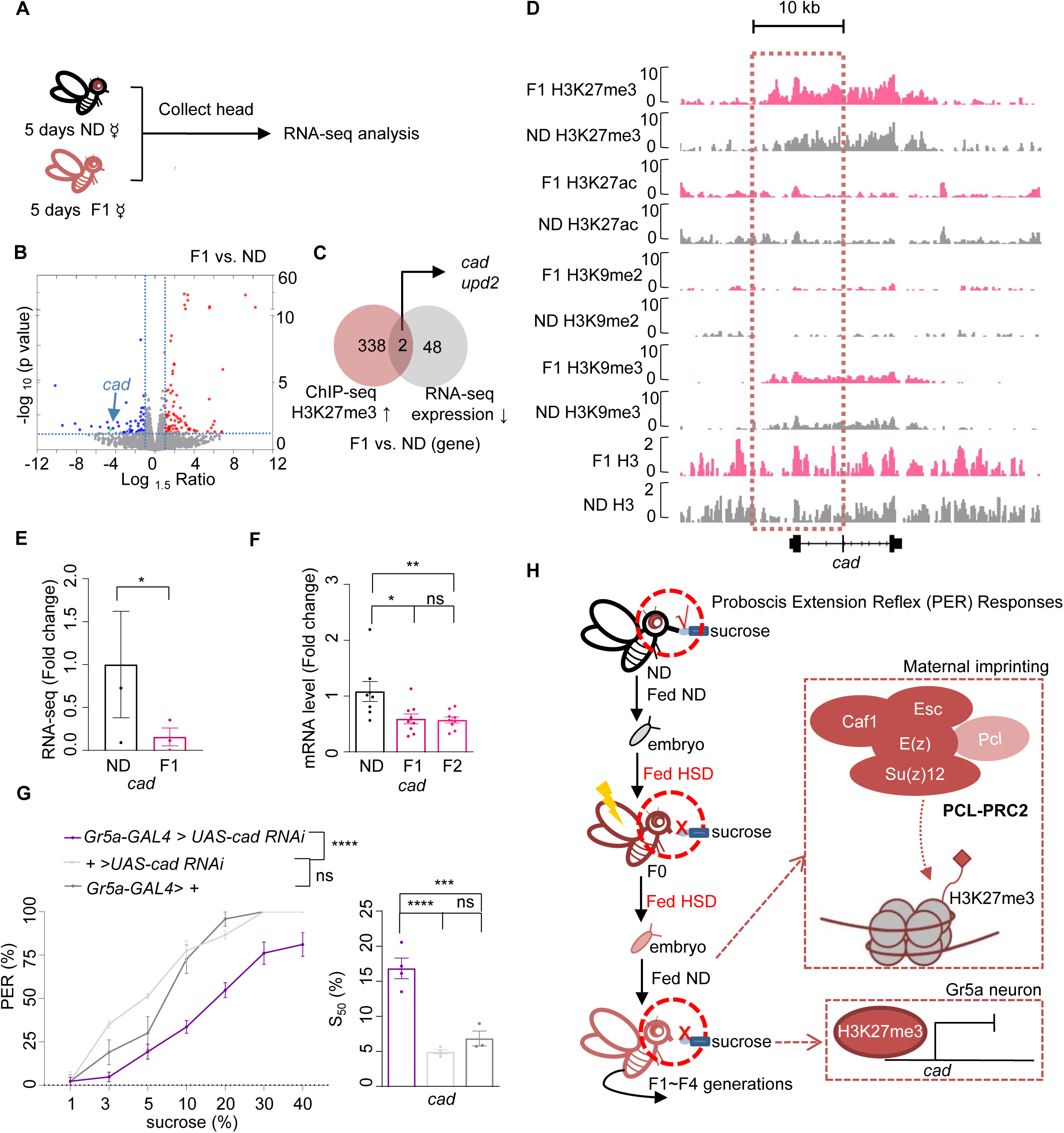
Cad mediated the inhibitory effect of H3K27me3 modifications on sweet sensitivity and feeding behavior. A. The preparation of RNA samples for RNA-seq. B. Volcano plot showed differentially expressed genes between ND and HSD-F1 flies. (blue: down-regulated genes in HSD-F1 flies; red: up-regulated genes in HSD-F1 flies; green: *cad*). The horizontal line indicated the significance threshold (p = 0.05) and the vertical lines indicated the 1.5-fold change threshold. C. Venn diagram of down-regulated genes and H3K27me3-target genes between ND and HSD-F1 flies (grey: down-regulated genes in the heads of HSD-F1 flies via RNA-seq analysis, fold change > 1.5; red: H3K27me3-target genes in HSD-F1 embryos via ChIP-seq analysis, log_10_ likelihood ratio>1). D. Genome browser view of H3K27me3, H3K27ac, H3K9me2, H3K9me3, and H3 density around the *cad* gene in embryos of ND and HSD-F1 flies. E-F. mRNA expression levels of *cad* in indicated flies. The fly heads were collected and subjected to RNA-seq (E) (n = 3 biological replicates, each containing 15 fly heads) or quantitative RT-PCR (F) (n = 8-9 biological replicates, each containing 15 fly heads). G. Fractions of flies of the indicated genotypes showing PER responses to different concentrations of sucrose (n = 3-6 biological replicates, each containing 8-12 flies). The S_50_ indicated the sucrose concentration that induced PER responses in 50% of the tested flies. H. A working model: Upon ancestral HSD exposure, PCL-PRC2 complex was activated and maintained high levels of H3K27me3 imprinting in maternal germline and the offspring. H3K27me3 targeted *cad* promoter regions and suppressed its expression, resulting in an inhibitory effect in Gr5a^+^ sweet-sensing gustatory neurons. Consequently, both sweet sensitivity and feeding behavior in the offspring were suppressed. Data are shown as means ± SEM. ns P>0.05; * P<0.05; ** P < 0.01; *** P < 0.001; **** P < 0.0001.

Among these genes, we noticed homeobox gene *caudal* (*cad*), a transcription factor involved in anterior/posterior patterning, organ morphogenesis, and innate immune homeostasis (Mlodzik & Gehring, 1987; Ryu et al., 2008), and *unpaired 2* (*upd2*), a *Drosophila* leptin ortholog and a secreted factor produced by the fat body and activated JAK/STAT signaling in GABAergic neurons (Hombría et al., 2005; Rajan & Perrimon, 2012), were the only two genes located in the H3K27me3 hypermethylated genome site and also showed down-regulated gene expression in HSD-F1 flies (*Figure 5B, C, and E, Figure 5-figure supplement 1C*). Given that Upd2 was mainly expressed in the fat body with clear metabolic functions (Brent & Rajan, 2020; Rajan & Perrimon, 2012), we focused on the ubiquitously expressed Cad for further behavioral analysis.

Notably, HSD-F1 flies exhibited a higher level of H3K27me3 modifications across the promoter region of *cad*, especially near the promoter region, whereas H3K9me3 signals exhibited no difference (*Figure 5D and Figure 5-figure supplement 1A, left)*. Other signals such as H3K27ac and H3K9me2 were generally weak around the promoter region of *cad* (*Figure 5D and Figure 5-figure supplement 1A*, *left*). We also conducted quantitative RT-PCR analysis and confirmed that Cad expression was down-regulated in both HSD-F1 and HSD-F2 flies (*Figure 5F*). These data indicate that ancestral HSD exposure elevates H3K27me3 levels in the promoter region of *cad* gene, resulting in a reduction in its mRNA expression in the head tissues of offspring.

We then asked whether Cad played a role in regulating sweet sensitivity in Gr5a^+^ gustatory neurons. Compared to the transgenic controls, knocking down Cad expression in Gr5a^+^ neurons led to lower PER responses to sucrose (*Figure 5G*). These results were consistent with a recent study and confirmed that Cad played an important role in regulating sweet sensitivity in Gr5a^+^ neurons (Vaziri et al., 2020). Furthermore, these results suggest that the reduction in Cad expression likely contributes to the deficits of sweet sensitivity and feeding behavior seen in the offspring of HSD-exposed ancestors.

It was unlikely that Cad was the sole gene mediated the transgenerational behavioral effect of HSD exposure. Other potential candidate genes might be missed out in the above analysis if they did not exhibit statistically different H3K27me3 modification levels or gene expression levels (*Figure 5B*). Besides Cad, we noticed several transcription factors, Ptx1, GATAe, nub, which were known to regulate sweet sensitivity (Vaziri et al., 2020), also exhibited elevated H3K27me3 modifications around their promoter regions. However, these genes didn’t show down-regulated gene expression in HSD-F1 flies (*Figure 5-figure supplement 2A* and B). It was thus possible that multiple transcriptional regulators were epigenetically modified by ancestral HSD exposure, which in turn exerted transgenerational effects on the sweet sensitivity and feeding behavior in offspring.

## DISCUSSION

Transgenerational behavioral change is present in many animal species. In *C. elegans*, exposure to pathogenic threats induces avoidance memories which can transmit for four generations via sRNA signaling (Moore et al., 2021). In fruit flies, exposure to predatory wasps leads to the inheritance of ethanol preference for five generations via maternal NPF repression (Bozler et al., 2019). Such a “behavior memory” may be evolutionarily beneficial in a sense to pre-adapt offspring for changing environmental conditions. Nevertheless, it may lead to devastating effects in human health.

Individuals who were exposed to Dutch famine in early gestation exhibited deficits in metabolism, cardiovascular health, and mental health (Schulz, 2010). Similarly, mice exposed to traumatic experiences and drugs induce depressive-like or autism-like behaviors in their progeny (Chan et al., 2020; Choi et al., 2016; Gapp et al., 2014).

Globally, HSD has become a routine of modern lifestyle, which is linked to various human diseases, including obesity, type 2 diabetes, and neurobiological diseases (Malik & Hu, 2022; Malik et al., 2010). HSD also induces many behavior disorders in animal models such as feeding abnormalities and addictive-like behaviors (Avena et al., 2009; May et al., 2019). This present study uncovers that HSD not only affects flies’ overall physiology and metabolism within the same generation, but also affects their sweet sensitivity and feeding behavior in a manner spanning multiple generations of offspring. If similar observations hold true in human society, HSD exposure may lead to an additional layer of health risk that needs to be recognized and addressed.

Mechanistically, our data indicate that elevated H3K27me3 modifications upon ancestral HSD exposure is the key epigenetic factor underlying the transgenerational regulations of sweet sensitivity and feeding behavior. HSD exposure enhances genome-wide H3K27me3 but not H3K27ac or H3K9me2/3 modifications in the early embryos. E(z) and Pcl, key components of PCL-PRC2 complex, play a critical role in catalyzing tri-methylation of the repressive chromatin marker histone H3 lysine 27 and maintaining such imprinting during fly development. Perturbation of PCL-PRC2 complex, via both genetic and pharmacological approaches, blocks the transmission of such repressive histone imprinting to offspring and eliminates the transgenerational modulation of sweet sensitivity and feeding behavior. Such epigenetic modulations are transmitted via the female germline, and exert a long-lasting effect on the expression of Cad (and possibly other transcriptional regulators) in the offspring. Cad, a transcription factor belonging to the Hox family, regulates the sensitivity of sweet-sensing gustatory neurons and plays a role in modulating PER responses to sucrose.

Feeding is tightly regulated by many factors, such as internal nutritional needs, overall physiological status, and environmental cues. However, most of these regulations are quite dynamic in nature and do not last for long. For example, in fruit flies energy shortage can trigger foraging and feeding behavior in the time scale of hours, which can be rapidly suppressed upon acquisition of desirable food sources (Basiri & Stuber, 2016; Qi et al., 2021). These regulations are often mediated by rapid-acting molecular and cellular mechanisms such as iron channels, hormones, and neuropeptides (Pool & Scott, 2014). HSD can also modulate feeding behavior in a dynamic manner (Sung et al., 2022). Moreover, this mechanism is reversible within the same generation. For example, the number of PLCβ2^+^ taste bud cells in the fungiform papilla decreased within 4 weeks of HSD exposure in mice and was completely restored within 4 weeks upon the removal of HSD (Sung et al., 2022). However, the mechanism in our present study is highly persistent and heritable. The deficits in sweet taste perception and feeding behavior can transmit through multiple generations without re-exposure to the original HSD treatment. These findings suggest that feeding behavior can be modulated by different regulatory mechanisms with distinct time scales in response to various types of environmental and internal state changes. The relationship between these different mechanisms will be of great interest to further study. For example, it remains unclear whether and how hormonal and neurotransmitter changes that occur upon HSD exposure in flies within the same generation contribute to the reprogramming of H3K27me3 that can last for multiple generations.

Based on our present study, several important questions remain to be answered. The first question is how HSD evaluates H3K27me3 modifications. Previous works reported that some dietary bioactive compounds could regulate histone modifying enzymes (Vahid et al., 2015). Thus, HSD may regulate the activity of histone methyltransferases (HMTs) or histone demethylases (HDMTs) directly. Alternatively, HSD may affect H3K27me3 modifications via certain nutrient-sensing mechanisms such as O-GlyNAc transferase, which is known to facilitate H3K27me3 formation with PRC2. The second question lies in how elevated H3K27me3 modifications in specific gene loci are retained during gamete formation and embryo development, and how such epigenetic imprinting is removed after 4-5 generations on ND. Previous work reported that long-term memory of H3K27me3 depended on efficient copying of this mark after each DNA replication cycle in a PRC2 dependent manner (Coleman & Struhl, 2017). It is therefore possible that PRC2-mediated maintenance of H3K27me3 imprinting can be regulated by dietary exposure. The third question is how H3K27me3 imprinting affects specific neurons in the adult via Cad signaling. We showed that up-regulated H3K27me3 modifications decreased Cad expression. Future studies are needed to examine how Cad signaling impacts the development and function of Gr5a^+^ gustatory neurons, and whether other transcription factors may also be involved. Previous work reported that Cad regulated a network containing 119 candidate genes that were implicated in sensory perception of chemical stimulus, neuropeptide signaling pathway, signal transduction, and transcription factor activity, which could impact the sweet gustatory neurons and play a role in modulating PER responses to sucrose (Vaziri et al., 2020).

It will also be of interest to explore other epigenetic mechanisms that may participate in the transgenerational inheritance of sweet perception, such as ncRNA and other histone modifications that have not been identified in this study. Especially, as the Piwi protein negatively regulates H3K27 trimethylation (Peng et al., 2016), future studies are needed to understand whether piRNAs that were reported direct transient heterochromatin formation and stabilize maternal mRNAs during embryogenesis (Dufourt et al., 2017; Fabry et al., 2021; Wang & Lin, 2021) are involved in the transgenerational inheritance we identified. Furthermore, the identification of HSD exposure-induced transgenerational sweet perception decrease in fruit flies raises the question of whether similar phenomenon and mechanism can be extended to other behaviors. Our findings highlight a novel and pivotal role of epigenetic modifications in preparing animals for the dynamic environment, which opens a new avenue of research to further uncover the interactions among prior experience, epigenetics, and behavioral modulations across generations.

## MATERIALS AND METHODS

### Flies

Flies were kept in vials containing a standard medium made of yeast, corn, and agar at 25°C, 60% relative humidity, and on a 12-hrs light-12-hrs dark cycle. Virgin female flies were collected shortly after eclosion and kept in groups (25 flies per vial) on standard fly medium (ND, with 10% sucrose) or HSD (ND plus additional 10% sucrose) for 4-6 days before experiments.

Fly strains used in the manuscript: *nosNGT-GAL4:* (*#*31777), *Maternal-tubulin-Gal4* (*#*2318), *Gr5a-GAL4* (#57592), and *UAS-GCaMP6m* (#42748) were obtained from the Bloomington *Drosophila* Stock Center at Indiana University; *UAS-E(z) RNAi* (#2831), *UAS-Pcl RNAi* (#1185), *UAS-Su(var)3-9 RNAi* (#3558), *UAS-Rpd3 RNAi* (#0695), and *UAS-cad RNAi* (#03877.N) were from the Tsinghua Fly Center.

### Chemicals and Antibodies

If not otherwise indicated, all chemicals were from Sigma. EED226 (5 μM, Selleck, USA, S8496), Chaetocine (5 μM, Selleck S8068), and A-395 (10 μM, Sigma-Aldrich, SML1923) were added to the standard medium. Flies were kept in these foods for 5 days before the assay (change fresh medium every two days).

The following antibodies were used: The antibodies against Histone H3 (di methyl K9) (ab1220), acetyl histone-h3-k27 (ab4729), Histone H3 (tri methyl K27) (ab6002), Histone H3 (tri methyl K9) (ab8898), and Histone H3 antibody (ab1791) were all purchased from Abcam.

### Triglyceride, glycogen, and trehalose measurement

For triglyceride and glycogen, single fly was anesthetized and transferred to tube with the corresponding extract. The samples were measured according to the manufacturer’s instructions. Glycogen were measured with Liver / Muscle glycogen assay kit (A043-1-1, Nanjing Jiancheng Bioengineering Institute, China). Triglyceride was measured with Triglyceride Quantification Colorimetric/Fluorometric Kit (A110-1-1, Nanjing Jiancheng Bioengineering Institute).

The hemolymph trehalose was quantified by trehalose quantification kit (A149-1-1, Nanjing Jiancheng Bioengineering Institute). Briefly, forty flies were anesthetized and then pierced in the thorax with dissecting forceps. The pierced flies were then transferred to perforated tubes and centrifuged for 5 mins at 3000 × g at 4°C to collect the hemolymph. Afterwards, 0.6 μL hemolymph was removed quickly into a 200 μL tube with trehalose extract and vortexed for 2-3 mins. After 45 mins standing, the sample was centrifuged at 8000 × g for 10 mins. Then 175 μL supernatant was added to 700 μL reaction solution and boiled for 5 mins. After cooling, 250 μL sample was removed to a 96-well plate for the light absorption value measurement at 620 nm.

### The CAFE assay

As described previously (Ja William et al., 2007), 25 indicated virgin flies were collected upon eclosion and aged for four days. To construct the CAFE set up, one holes were bored into the lid of a *Drosophila* bottle and inserted a 5-μl glass capillaries (VWR, 53432-604) which filled with 20% sucrose. The bottle also contained 2% agar medium to ensure satiation with water. 10 female virgin flies were inserted into each bottle by mouth aspiration and adapted 24 hrs, then change new capillaries containing same concentration sucrose and the level of capillary was marked. After 24 hrs at 25°C, 60% relative humidity, the level of capillary was marked again and the distance between these marks converted into a volume consumed per fly. In addition, three blank bottles without flies were set up in the same way, and the mean volume change from these capillaries subtracted from the capillaries with flies, to control for the effect of evaporation.

### The MAFE assay

As described previously (Qi et al., 2015), individual flies were transferred and immobilized in a 200 μL pipette tip, and then sated with sterile water before being presented with 400 mM sucrose filled in a graduated glass capillary (VWR, #53432-604). The food stimulation was repeated until the flies became unresponsive to a series of ten food stimuli, and the total food consumption was calculated based on the volume change during feeding process

### Proboscis extension response (PER) assay

PER assay was performed as described (Qi et al., 2015). Briefly, individual flies were gently aspirated and immobilized in a 200 μL pipette tip as the MAFE assay. Flies were first sated with water and then subjected to different sugar solutions with each solution tested twice. Flies showing PER responses to at least one of the two trials were considered positive to that sugar concentration. All PER experiments are n > 3 replicates with 8-12 flies per replicate unless otherwise stated.

The S_50_ indicated the sucrose concentration that induces 50 % PER, which was estimated using the basic linear or nonlinear regression model based on a previous method (Vaziri et al., 2020). All S_50_ estimations were performed in R package using the “basicTrendline” function.

### Quantitative RT-PCR

Total RNA was extracted from the head tissues of flies. RNA was reverse transcribed with the All-in-one cDNA synthesis supermix (TransScript). Quantitative RT-PCR was conducted on Bio-Rad CFX96 using the SYBR green PCR master mix (TaKaRa, Japan) with the primers listed in Appendix 1—key resources table. Relative mRNA levels were calculated relative to *rp49* expression by the comparative Ct method.

### Calcium imaging

For *in vivo* imaging (Yang et al., 2018), flies were anesthetized on ice and glued onto transparent tape. Then a hole (∼1–2 mm) on the tape was incised to expose the dorsal part of the fly head. The cuticle part around the Gr5a^+^ neurons region of the fly brain was gently removed with forceps and the brain was bathed in the adult hemolymph-like solution (AHL; 108 mM NaCl, 8.2 mM MgCl_2_, 4 mM NaHCO_3_, 1 mM NaH2PO_4_, 2 mM CaCl_2_, 5 mM KCl, 5 mM HEPES, 80 mM sucrose, pH7.3). A micro manipulator delivered liquid food to the proboscis of the fly at the indicated time and the actual feeding bouts were imaged by a digital camera installed under the imaging stage at 0.5 frame/s.

More specifically, at each feeding bout, the flies extended their proboscis to reach the surface of the liquid food and started food ingestion. By adding a blue dye in the liquid food, the actual flow of the dyed food through flies’ pharynx could also be seen.

The calcium signals of Gr5a^+^ neurons were recorded by a Nikon C2 confocal microscope, with a water immersion objective lens (40× /0.80 w DIC N2) at 0.2 frame/s. Image analyses were performed in ImageJ and plotted in Excel (Microsoft). The ratio changes were calculated using the following formula: ΔF/F = [F – F_0_]/F_0_, where F is the mean florescence of cell body, F_0_ is the average base line (15∼30s interval before stimulation).

### RNA-seq and analysis

Total RNA from fly heads was extracted from 5-day-old female flies using the Trizol reagent (Invitrogen, USA), mRNA was purified from total RNA using oligo(dT)-attached magnetic beads, followed by library preparation [The quality of libraries was checked by Bioanalyzer 2100 (Agilent)] and sequencing (BGISEQ 500 platform) with paired end 150bps. Sequence data were subsequently mapped to *Drosophila* genome and uniquely mapped reads were collected or further analysis. Gene expression was calculated by the FPKM (Fragments Per Kilobase Of Exon Per Million Fragments Mapped). The genes with p-value less 0.05 and |log_2_FoldChange| more than 1 were considered as the differentially expressed gene. The RNA-seq data were deposited in GEO database under the accession code (GSE216075 and GSE215756).

### Embryo sorting

Flies were maintained in large population cages in an incubator set at standard conditions (25°C). Embryos were collected for 30 mins, and then allowed to develop for 50 additional minutes before being harvested. Harvested embryos were preserved in the embryo stock buffer A1 (60 mM KCl, 15 mM NaCl, 4 mM MgCl_2_, 15 mM HEPES, 0.5% Triton X-100, 0.5 mM DTT and 10 mM sodium butyrate) and hand sorted (within 30 mins) in small dish using an inverted microscope to remove embryos younger or older than the targeted age range based on morphology of the embryos as previously described (Li et al., 2014).

### ChIP-seq and analysis

Formaldehyde was added to the embryos for cross-linking for 10 mins, then glycine was added to quench the formaldehyde, followed by centrifugation and removal of the supernatant. The pellet was washed twice and then lysed with the Lysis Buffer (140mM NaCl, 15mM HEPES, 1mM EDTA, 0.5mM EGTA, 1% Triton X-100, 0.1% sodium deoxycholate). The lysate is sonicated using Qsonica (Duty Cycle--10%; Intensity--5; Cycles per Burst--200; Time--4mins) and centrifuged. The chromatin obtained was fragmented to sizes ranging from 100 to 300 bp. The supernatant was carefully transferred to a new tube to incubate with corresponding antibody (10 μg) at 4°C over night. Then 15 μL of Pierce™ Protein A/G beads (Thermo Fisher) were added to the mixture, followed by further incubation for 4 hrs on a rotator. After washing 4 times with TE Buffer (0.1mM EDTA 0.5M, 10mM Tris-HCl pH 8.0), the beads were incubated with the 250 μl Elution Buffer (10 mM EDTA, 50 mM Tris HCl pH 8.0, 1% SDS) at 65°C for 30 mins. Then 1.6 μl 25mg/ml RNase A (Sigma) were added, and all samples were incubated at 65°C for 3hrs. After the RNase A treatment, all sample were further treated with 6 μl Proteinase K (Sigma) at 56°C for 2hrs. The resulted DNA was purified by Agencourt Ampure beads (Beckman Coulter). 1 μg of DNA was used to generate sequencing library using the mRNA-Seq Sample Preparation Kit (Illumina) and sequenced on an Illumina Hiseq platform (Novagene) with paired end 300 bps.

Raw reads were cleaned using trim galore. The reads were then aligned to the dm6 genome assembly using Bowtie2 v2.3.5.1 with default parameters. Duplicate reads were then removed using MarkDuplicates from gatk package v.4.1.4.1. Replicate samples were merged using the samtools v1.10. For ChIP-seq, bigwig tracks were generate using bamCompare from deeptools 3.3.1 (parameters: --skipNAs -- scaleFactorsMethod CPM --operation log2 --extendReads 200). Negative values were set to zero. Peak calling was performed using Macs2 v2.2.6 callpeak with default parameters. ChIP-seq profiles were created by computeMatrix and plotProfile in deeptools 3.3.1. IGV v.2.4.13 was used to visualize the bigwig tracks. The difference peaks of ChIP-seq data were found by using MACS2 with the options “bdgdiff”. The parameters are “--t1 --c1 --t2 --c2 --d1 --d2 --o-prefix” and others were default.

### Statistical analysis

Data presented in this study were verified for normal distribution by D’Agostino– Pearson omnibus test. Student’s t test, one-way ANOVA, and two-way ANOVA (for comparisons among three or more groups and comparisons with more than one variant) were used. The post hoc test with Bonferroni correction was performed for multiple comparisons following ANOVA.

## ACKNOWLEDGEMENTS

We thank all Wang Lab members for helpful discussions and technical assistance. We thank Dr. Wei Xie from Tsinghua University and Neuroscience Pioneer Club for helpful discussions throughout the course of this study. Ye Wu and Tingting Song provided scientific and administrative support in the laboratory. We thank the Bloomington *Drosophila* stock center at Indiana University and the Tsinghua Fly Center for fly stocks and reagents. This study was funded by National Key R&D Program of China (2019YFA0802400 and 2019YFA0801900), the National Natural Science Foundation of China (32071006), and the startup funds from Shenzhen Bay Laboratory.

---

**Figure 1-figure supplement 1.**
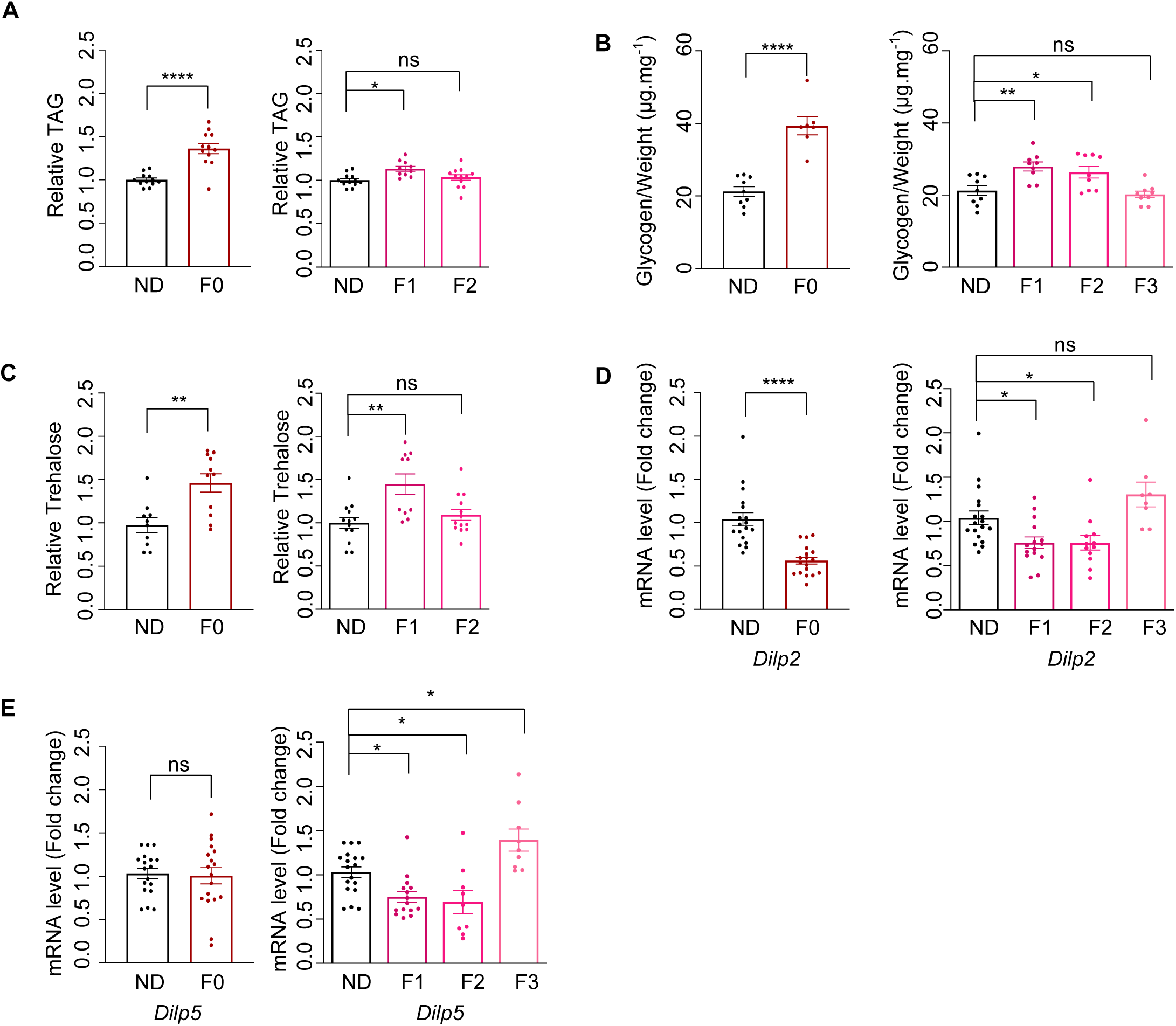
Ancestral HSD exposure induced metabolic changes in the offspring. A-B. The levels of triglyceride (A), glycogen (B) and trehalose (C) of individual female flies of the indicated groups (n = 8-15 biological replicates, each containing 5 flies). D-E. mRNA expression levels of *Dilp2* (D) and *Dilp5* (E) in the head tissues of indicated flies by qPCR (n = 8-15 biological replicates, each containing 3 flies).

**Figure 1-figure supplement 2.**
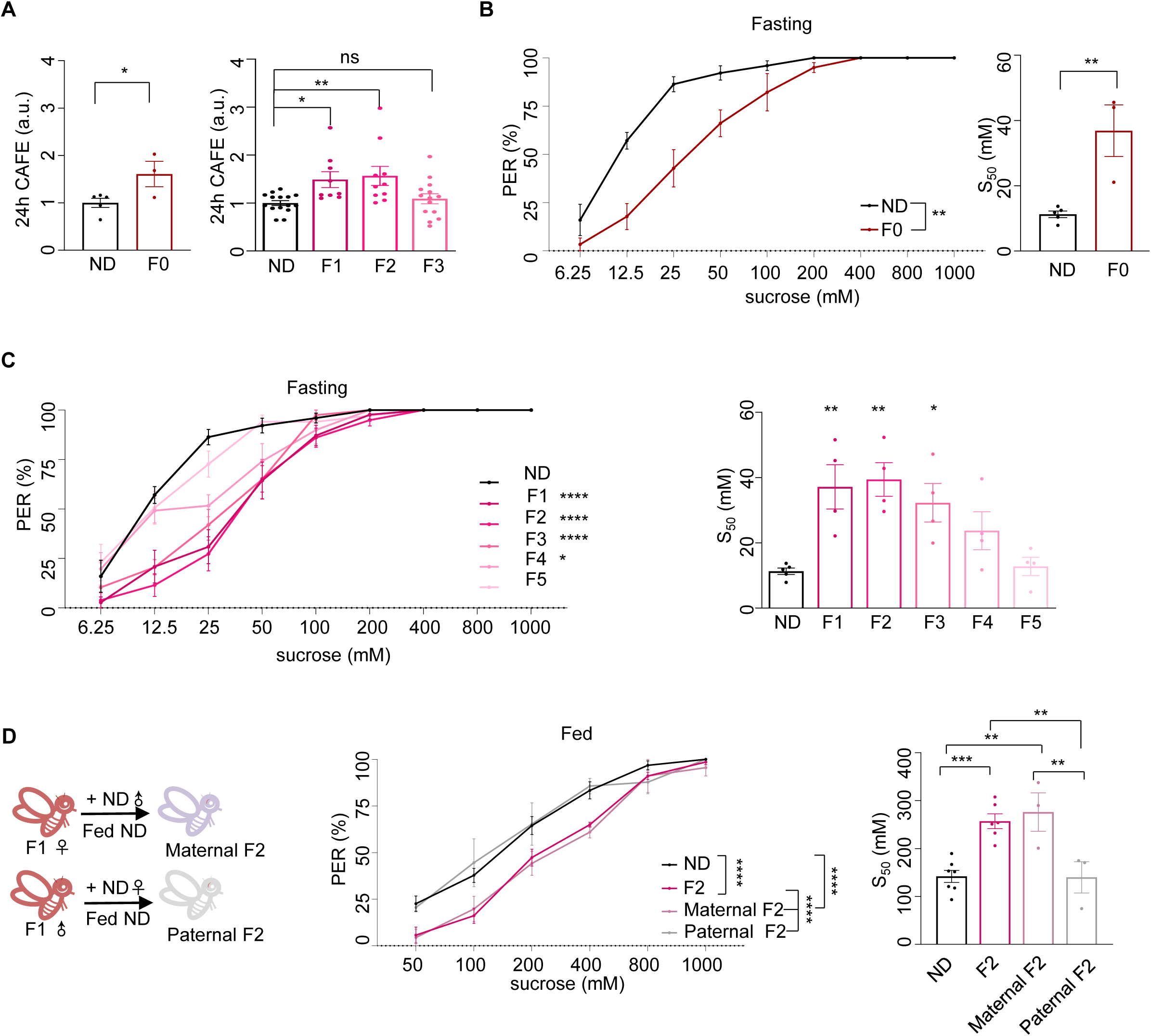
Ancestral HSD exposure induced behavioral changes in the offspring. A. Volume of 20% sucrose consumed by indicated flies during 24 hrs using the CAFE assay (Normalized with ND flies, n = 3-15 biological replicates, each containing 20 flies). B-D. Fractions of indicated flies showing PER responses to different concentrations of sucrose (n = 3-6 biological replicates, each containing 8-12 flies). The S_50_ indicated the sucrose concentration that induced PER responses in 50% of the tested flies. The flies in B and C were pre-starved for 12 hrs (fasting) before the assay. Data were shown as means ± SEM. ns P>0.05; * P<0.05; ** P < 0.01; *** P < 0.001; **** P < 0.0001.

**Figure 2-figure supplement 1.**
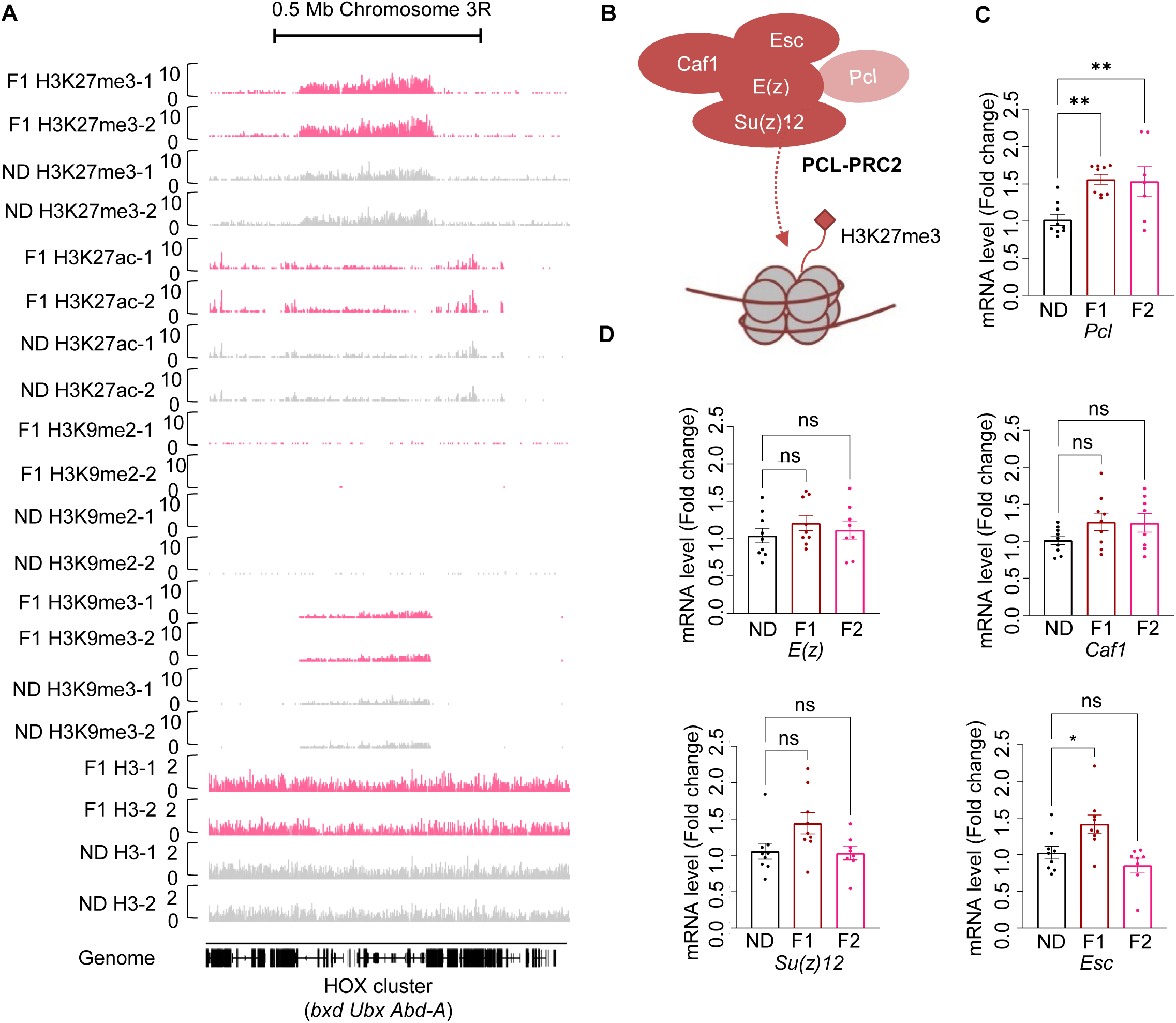
Ancestral HSD exposure enhanced *Pcl* expression in the offspring. A. Genome browser view of H3K27me3, H3K27ac, H3K9me2, and H3K9me3 density at the HOX cluster (bxd, Ubx, and Abd-A) gene regions in embryos of two replicates ND and F1 flies. B. Schematic diagram of PCL-PRC2 complex. Polycomb repressive complex 2 (PRC2) is a chromatin-modifying enzyme that catalyzes the methylation of histone H3 at lysine 27 and contains the Enhancer of zeste [E(z), the enzymatic component of PRC2, Suppressor of zeste 12 [Su(z)12], Extra sex combs (Esc), and Chromatin assembly factor-1 (Caf-1). Polycomb-like (PCL) interacts with PRC2 to constitute a specific form of PCL-PRC2 complex to generate high levels of H3K27me3. C-D. mRNA expression levels of *Pcl* (C), *E(z)*, *Caf-1*, *Su(z)12*, and *Esc* (D) of indicated flies (n = 7-9 biological replicates, each containing 3 flies). Data were shown as means ± SEM. ns P>0.05; * P<0.05; ** P < 0.01; *** P < 0.001; **** P < 0.0001.

**Figure 3-figure supplement 1.**
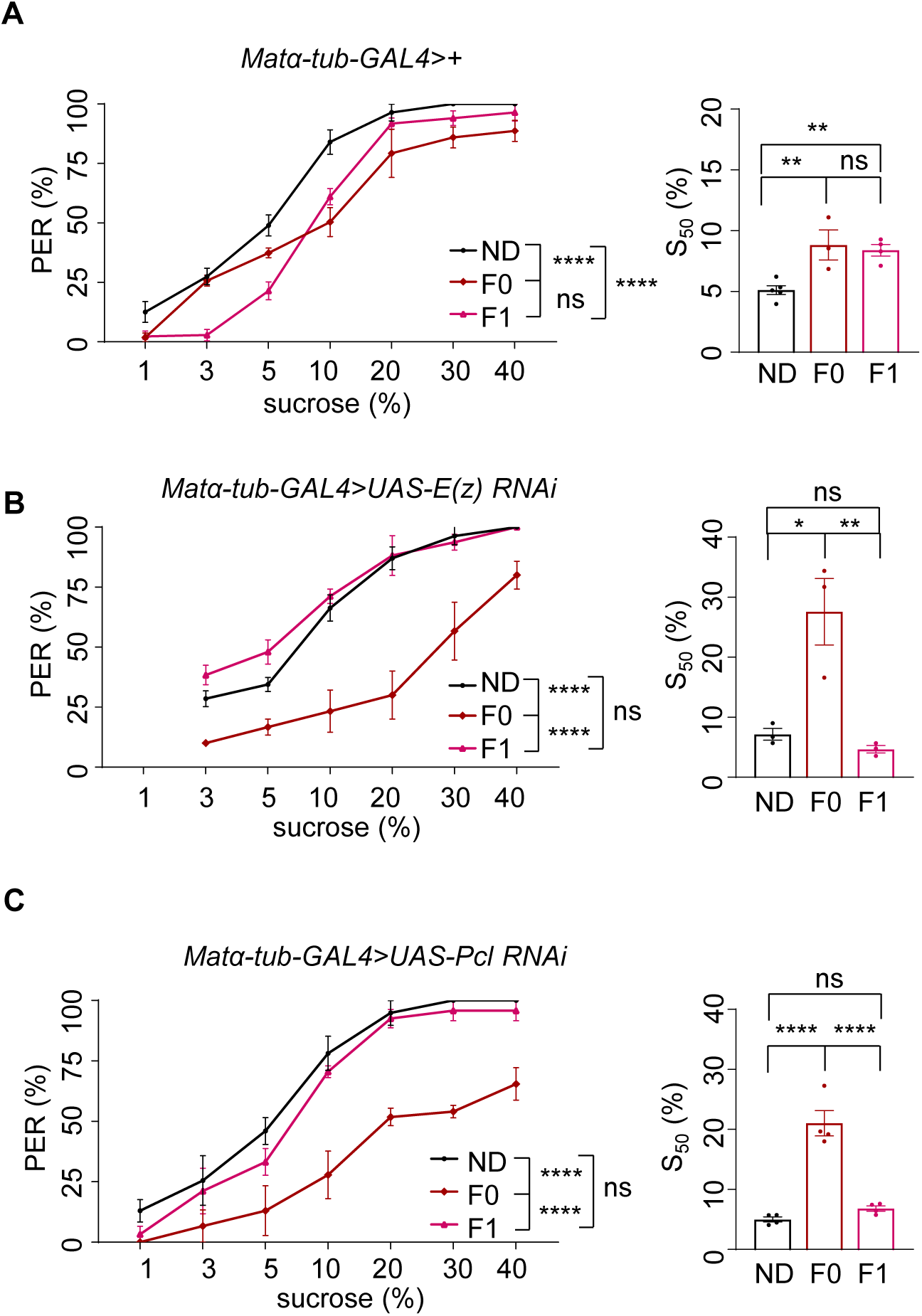
H3K27me3 in the female germline was required for transgenerational modulation of sweet sensitivity and feeding behavior. A-C. Fractions of flies of the indicated genotypes showing PER responses to different concentrations of sucrose (n = 3-6 biological replicates, each containing 8-12 flies). The S_50_ indicated the sucrose concentration that induced PER responses in 50% of the tested flies. Data were shown as means ± SEM. ns P>0.05; * P<0.05; ** P < 0.01; *** P < 0.001; **** P < 0.0001.

**Figure 5-figure supplement 1.**
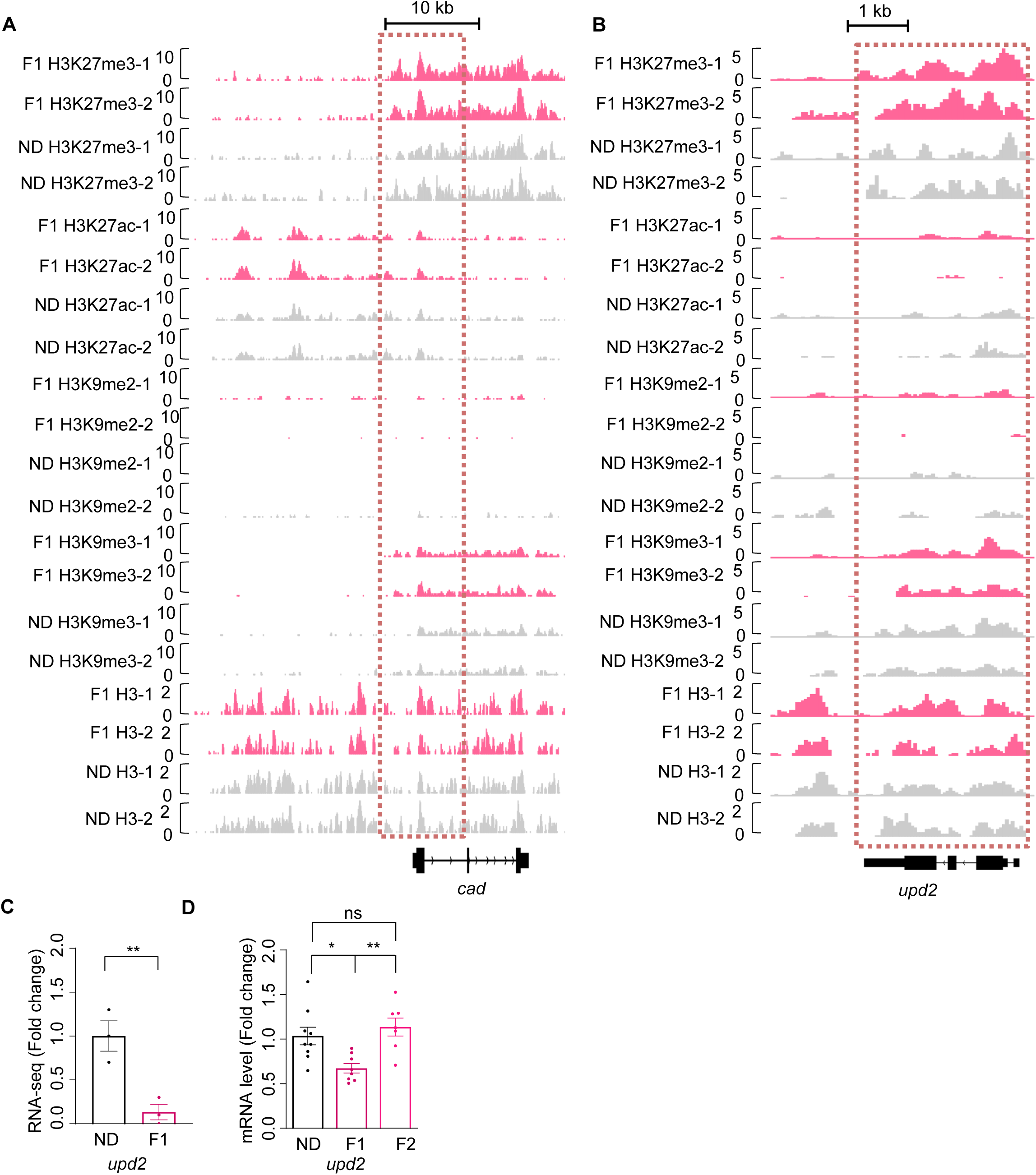
H3K27me3 modifications and mRNA expression of *cad* and *upd2* genes upon ancestral HSD exposure. A-B. Genome browser view of H3K27me3, H3K27ac, H3K9me2, H3K9me3, and H3 density at the *cad* (A) and *upd2* (B) genes in embryos of ND and HSD-F1 flies. C-D. mRNA expression levels of *upd2* in indicated flies. The fly heads were collected and subjected to RNA-seq (C) (n = 3 biological replicates, each containing 15 fly heads) or quantitative RT-PCR (D) (n = 6-9 biological replicates, each containing 3 flies). Data were shown as means ± SEM. ns P>0.05.

**Figure 5-figure supplement 2.**
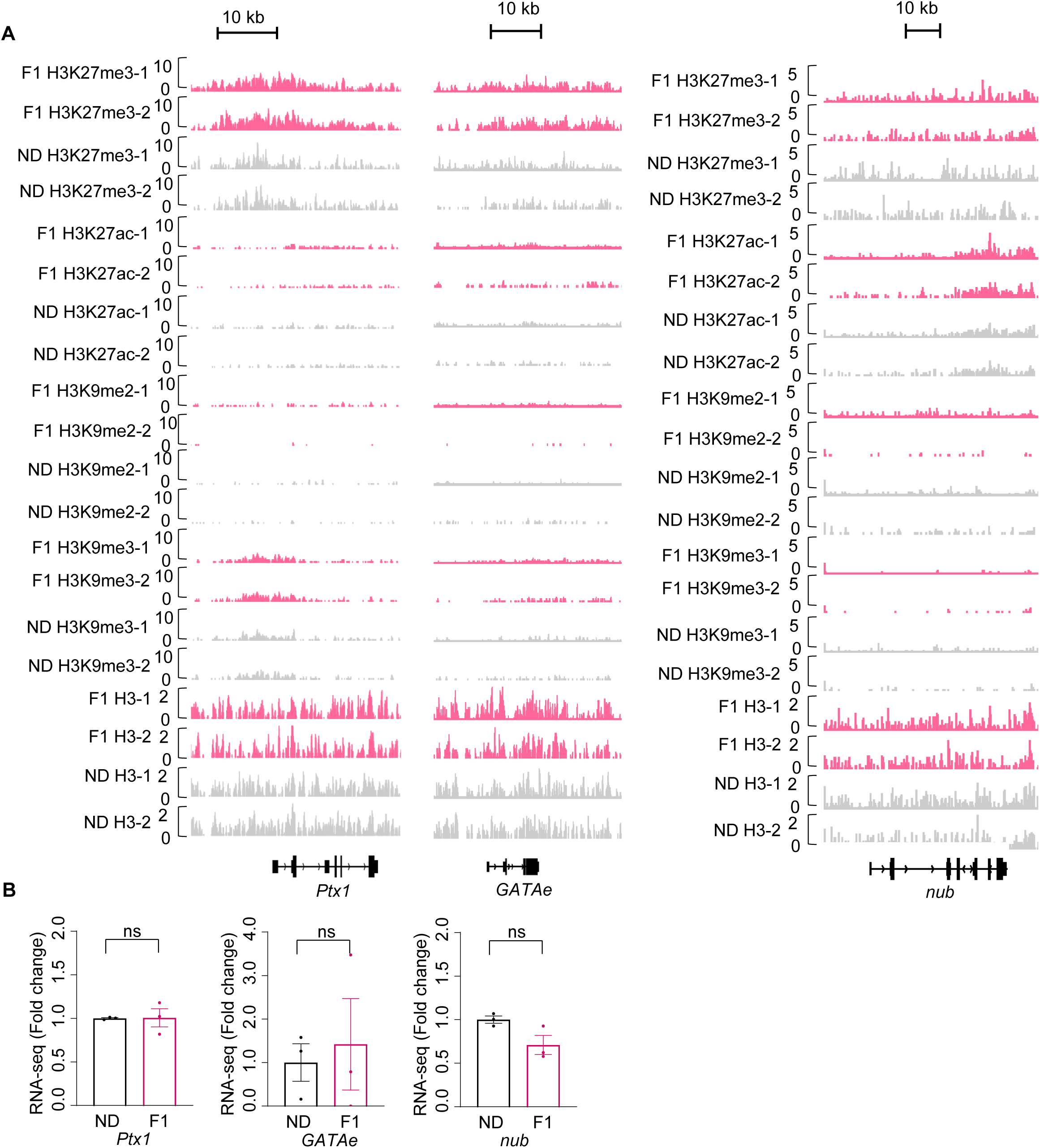
H3K27me3 modifications and mRNA expression of *Ptx1, GATAe and nub* upon ancestral HSD exposure. A. Genome browser view of H3K27me3, H3K27ac, H3K9me2, H3K9me3, and H3 density at the *Ptx1* (*left*), *GATAe* (*middle*) and *nub* (*right*) genes in embryos of ND and HSD-F1 flies. B. mRNA expression levels of *Ptx1*, *GATAe*, and *nub* in indicated flies. The fly heads were collected and subjected to RNA-seq (n = 3 biological replicates, each containing 15 fly heads). Data were shown as means ± SEM. ns P>0.05.

## Appendix 1

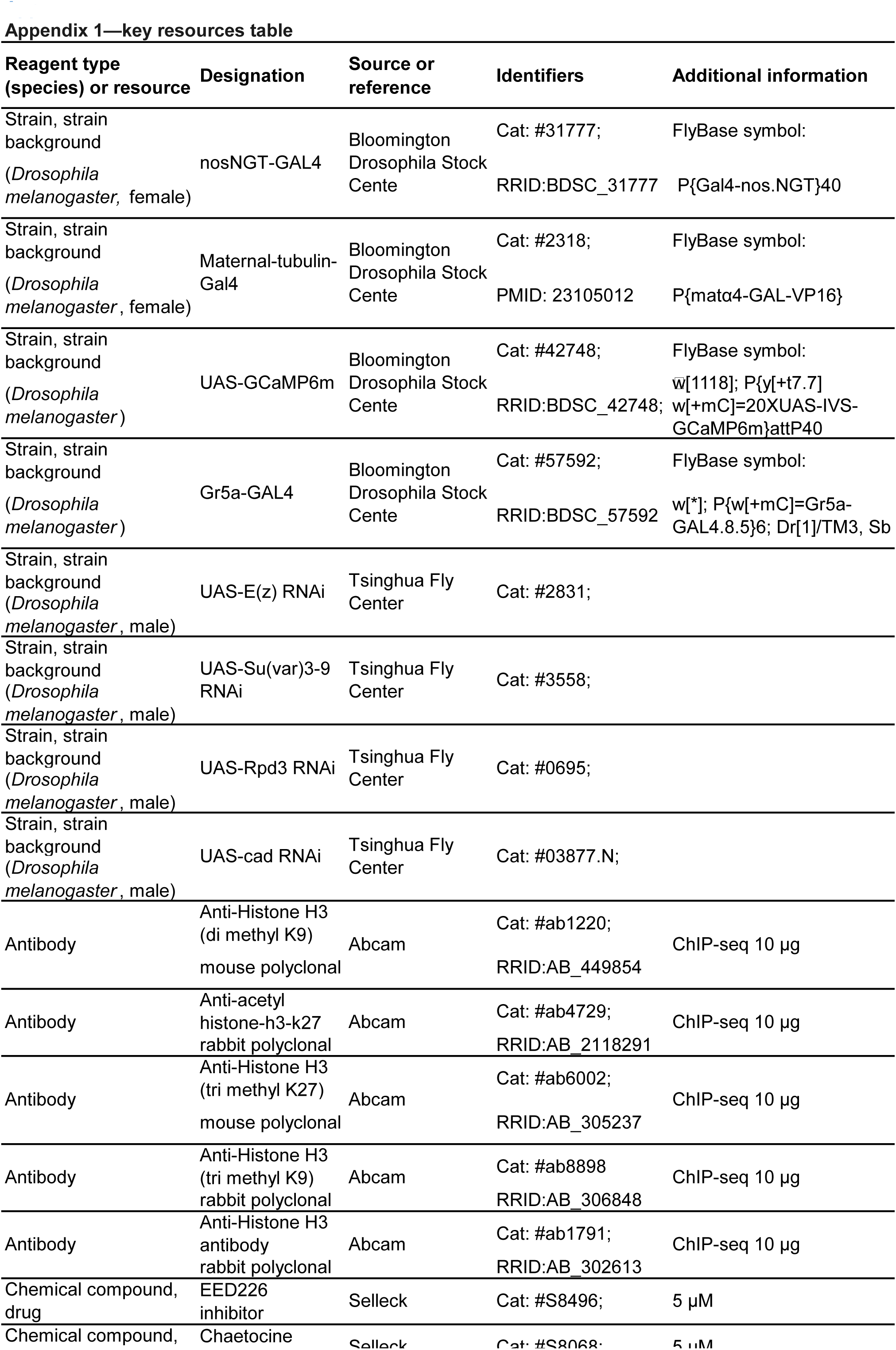
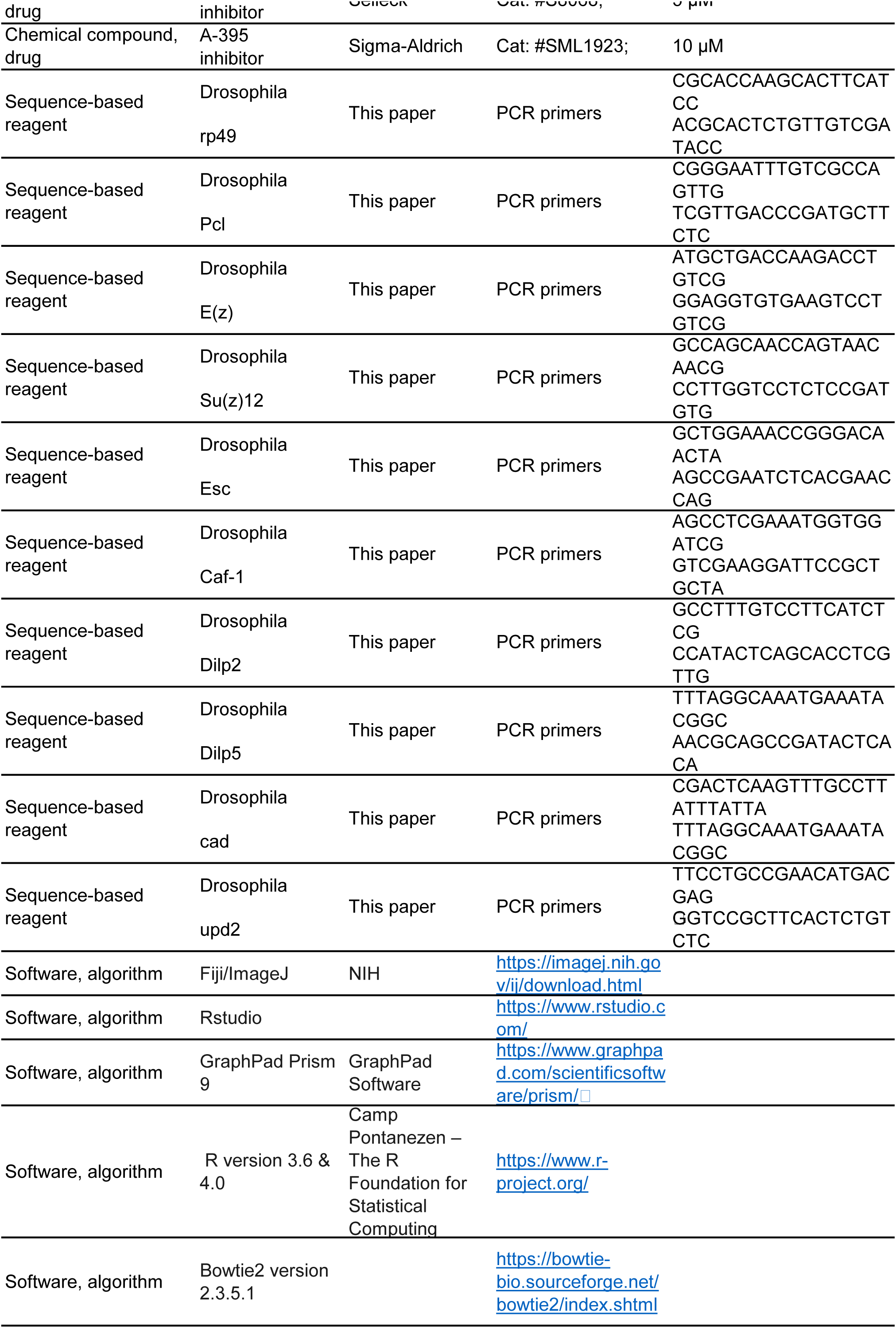
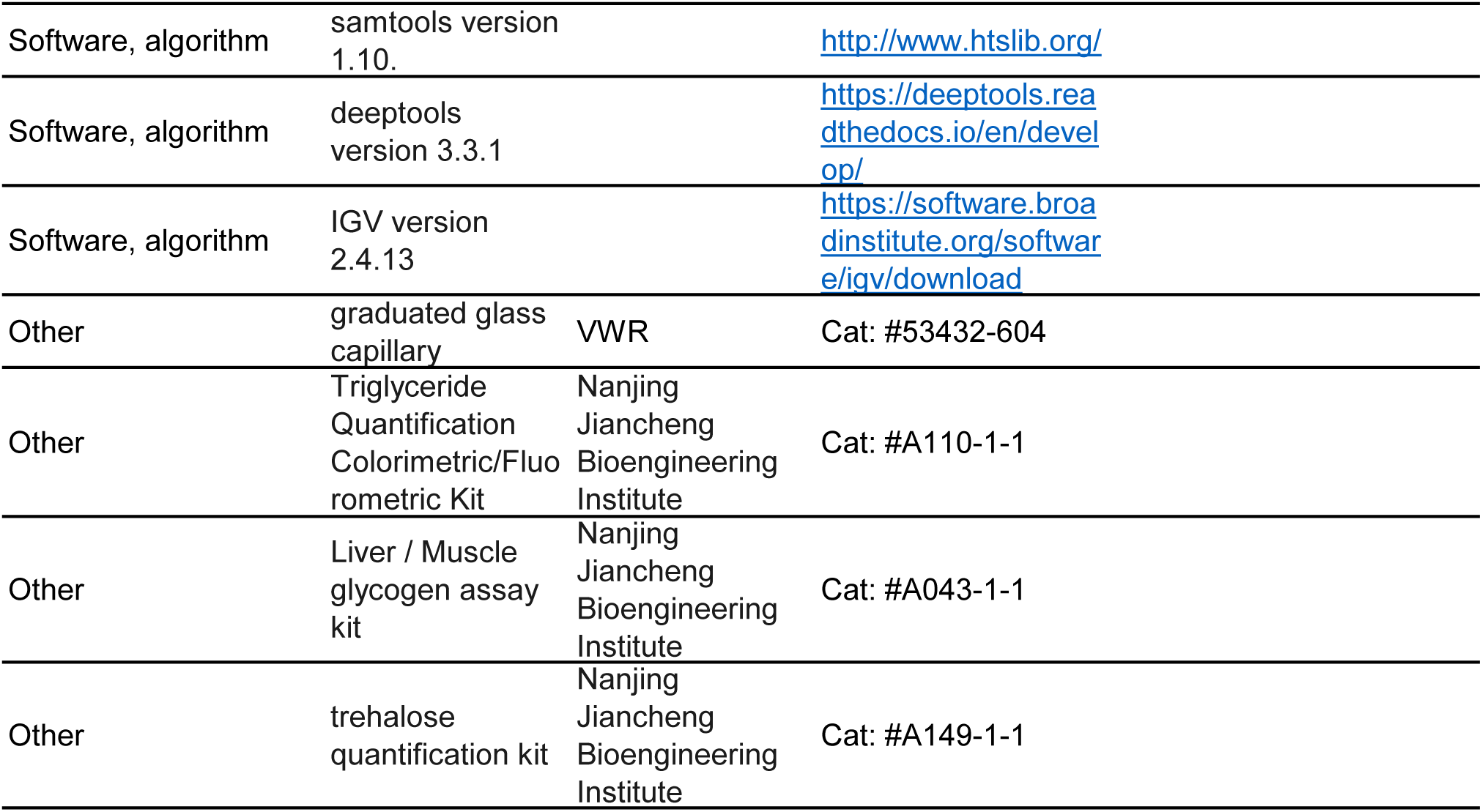

## REFERENCES

1. Aldrich, J. C., & Maggert, K. A. (2015). Transgenerational Inheritance of Diet-Induced Genome Rearrangements in Drosophila. PLOS Genetics, 11(4), e1005148. https://doi.org/10.1371/journal.pgen.1005148

2. Anson, R. M., Guo, Z., de Cabo, R., Iyun, T., Rios, M., Hagepanos, A., Ingram, D. K., Lane, M. A., & Mattson, M. P. (2003). Intermittent fasting dissociates beneficial effects of dietary restriction on glucose metabolism and neuronal resistance to injury from calorie intake. Proceedings of the National Academy of Sciences, 100(10), 6216–6220. https://doi.org/10.1073/pnas.1035720100

3. Arnold, S. E., Lucki, I., Brookshire, B. R., Carlson, G. C., Browne, C. A., Kazi, H., Bang, S., Choi, B.-R., Chen, Y., McMullen, M. F., & Kim, S. F. (2014). High fat diet produces brain insulin resistance, synaptodendritic abnormalities and altered behavior in mice. Neurobiology of Disease, 67, 79–87. https://doi.org/10.1016/j.nbd.2014.03.011

4. Avena, N. M., Rada, P., & Hoebel, B. G. (2009). Sugar and Fat Bingeing Have Notable Differences in Addictive-like Behavior. The Journal of Nutrition, 139(3), 623–628. https://doi.org/10.3945/jn.108.097584

5. Basiri, M. L., & Stuber, G. D. (2016). Multimodal Signal Integration for Feeding Control. Cell, 165(3), 522–523. https://doi.org/10.1016/j.cell.2016.04.022

6. Birse, R. T., Choi, J., Reardon, K., Rodriguez, J., Graham, S., Diop, S., Ocorr, K., Bodmer, R., & Oldham, S. (2010). High-fat-diet-induced obesity and heart dysfunction are regulated by the TOR pathway in Drosophila. Cell Metab, 12(5), 533–544. https://doi.org/10.1016/j.cmet.2010.09.014

7. Bohacek, J., & Mansuy, I. M. (2015). Molecular insights into transgenerational non-genetic inheritance of acquired behaviours. Nature Reviews Genetics, 16(11), 641–652. https://doi.org/10.1038/nrg3964

8. Bozler, J., Kacsoh, B. Z., & Bosco, G. (2019). Transgenerational inheritance of ethanol preference is caused by maternal NPF repression. eLife, 8, e45391. https://doi.org/10.7554/eLife.45391

9. Brent, A. E., & Rajan, A. (2020). Insulin and Leptin/Upd2 Exert Opposing Influences on Synapse Number in Fat-Sensing Neurons. Cell Metabolism, 32(5), 786–800.e787. https://doi.org/10.1016/j.cmet.2020.08.017

10. Buescher, J. L., Musselman, L. P., Wilson, C. A., Lang, T., Keleher, M., Baranski, T. J., & Duncan, J. G. (2013). Evidence for transgenerational metabolic programming in rosophila. Disease Models & Mechanisms, 6(5), 1123–1132. https://doi.org/10.1242/dmm.011924

11. Buettner, R., Schölmerich, J., & Bollheimer, L. C. (2007). High-fat Diets: Modeling the Metabolic Disorders of Human Obesity in Rodents. Obesity, 15(4), 798–808. https://doi.org/10.1038/oby.2007.608

12. Chan, J. C., Morgan, C. P., Adrian Leu, N., Shetty, A., Cisse, Y. M., Nugent, B. M., Morrison, K. E., Jašarević, E., Huang, W., Kanyuch, N., Rodgers, A. B., Bhanu, N. V., Berger, D. S., Garcia, B. A., Ament, S., Kane, M., Neill Epperson, C., & Bale, T. L. (2020). Reproductive tract extracellular vesicles are sufficient to transmit intergenerational stress and program neurodevelopment. Nature Communications, 11(1), 1499. https://doi.org/10.1038/s41467-020-15305-w

13. Chen, B., Du, Y.-R., Zhu, H., Sun, M.-L., Wang, C., Cheng, Y., Pang, H., Ding, G., Gao, J., Tan, Y., Tong, X., Lv, P., Zhou, F., Zhan, Q., Xu, Z.-M., Wang, L., Luo, D., Ye, Y., Jin, L., … Huang, H. (2022). Maternal inheritance of glucose intolerance via oocyte TET3 insufficiency. Nature, 605(7911), 761–766. https://doi.org/10.1038/s41586-022-04756-4

14. Chen, D., Yang, J., Xiao, Z., Zhou, S., & Wang, L. (2021). A diet-induced type 2 diabetes model in Drosophila. Science China Life Sciences, 64(2), 326–329. https://doi.org/10.1007/s11427-020-1774-y

15. Chen, Q., Yan, M., Cao, Z., Li, X., Zhang, Y., Shi, J., Feng, G.-h., Peng, H., Zhang, X., Zhang, Y., Qian, J., Duan, E., Zhai, Q., & Zhou, Q. (2016). Sperm tsRNAs contribute to intergenerational inheritance of an acquired metabolic disorder. Science, 351(6271), 397–400. https://doi.org/10.1126/science.aad7977

16. Chng, W.-b. A., Hietakangas, V., & Lemaitre, B. (2017). Physiological Adaptations to Sugar Intake: New Paradigms from Drosophila melanogaster. Trends in Endocrinology & Metabolism, 28(2), 131–142. https://doi.org/10.1016/j.tem.2016.11.003

17. Choi, C. S., Gonzales, E. L., Kim, K. C., Yang, S. M., Kim, J.-W., Mabunga, D. F., Cheong, J. H., Han, S.-H., Bahn, G. H., & Shin, C. Y. (2016). The transgenerational inheritance of autism-like phenotypes in mice exposed to valproic acid during pregnancy. Scientific Reports, 6(1), 36250. https://doi.org/10.1038/srep36250

18. Coleman, R. T., & Struhl, G. (2017). Causal role for inheritance of H3K27me3 in maintaining the OFF state of a Drosophila HOX gene. Science, 356(6333). https://doi.org/10.1126/science.aai8236

19. Dahanukar, A., Lei, Y.-T., Kwon, J. Y., & Carlson, J. R. (2007). Two Gr Genes Underlie Sugar Reception in Drosophila. Neuron, 56(3), 503–516. https://doi.org/10.1016/j.neuron.2007.10.024

20. Daxinger, L., & Whitelaw, E. (2012). Understanding transgenerational epigenetic inheritance via the gametes in mammals. Nat Rev Genet, 13(3), 153–162. https://doi.org/10.1038/nrg3188

21. Dufourt, J., Bontonou, G., Chartier, A., Jahan, C., Meunier, A.-C., Pierson, S., Harrison, P. F., Papin, C., Beilharz, T. H., & Simonelig, M. (2017). piRNAs and Aubergine cooperate with Wispy poly(A) polymerase to stabilize mRNAs in the germ plasm. Nature Communications, 8(1), 1305. https://doi.org/10.1038/s41467-017-01431-5

22. Dunn, G. A., & Bale, T. L. (2011). Maternal High-Fat Diet Effects on Third-Generation Female Body Size via the Paternal Lineage. Endocrinology, 152(6), 2228–2236. https://doi.org/10.1210/en.2010-1461

23. Fabry, M. H., Falconio, F. A., Joud, F., Lythgoe, E. K., Czech, B., & Hannon, G. J. (2021). Maternally inherited piRNAs direct transient heterochromatin formation at active transposons during early Drosophila embryogenesis. eLife, 10, e68573. https://doi.org/10.7554/eLife.68573

24. Gapp, K., Jawaid, A., Sarkies, P., Bohacek, J., Pelczar, P., Prados, J., Farinelli, L., Miska, E., & Mansuy, I. M. (2014). Implication of sperm RNAs in transgenerational inheritance of the effects of early trauma in mice. Nat Neurosci, 17(5), 667–669. https://doi.org/10.1038/nn.3695

25. Guida, M. C., Birse, R. T., Dall’Agnese, A., Toto, P. C., Diop, S. B., Mai, A., Adams, P. D., Puri, P. L., & Bodmer, R. (2019). Intergenerational inheritance of high fat diet-induced cardiac lipotoxicity in Drosophila. Nature Communications, 10(1), 193. https://doi.org/10.1038/s41467-018-08128-3

26. He, Y., Korboukh, I., Jin, J., & Huang, J. (2012). Targeting protein lysine methylation and demethylation in cancers. Acta Biochimica et Biophysica Sinica, 44(1), 70–79. https://doi.org/10.1093/abbs/gmr109

27. He, Y., Selvaraju, S., Curtin, M. L., Jakob, C. G., Zhu, H., Comess, K. M., Shaw, B., The, J., Lima-Fernandes, E., Szewczyk, M. M., Cheng, D., Klinge, K. L., Li, H.-Q., Pliushchev, M., Algire, M. A., Maag, D., Guo, J., Dietrich, J., Panchal, S. C., … Pappano, W. N. (2017). The EED protein–protein interaction inhibitor A-395 inactivates the PRC2 complex. Nature Chemical Biology, 13(4), 389–395. https://doi.org/10.1038/nchembio.2306

28. Heard, E., & Martienssen, Robert A. (2014). Transgenerational Epigenetic Inheritance: Myths and Mechanisms. Cell, 157(1), 95–109. https://doi.org/10.1016/j.cell.2014.02.045

29. Heijmans, B. T., Tobi, E. W., Stein, A. D., Putter, H., Blauw, G. J., Susser, E. S., Slagboom, P. E., & Lumey, L. H. (2008). Persistent epigenetic differences associated with prenatal exposure to famine in humans. Proceedings of the National Academy of Sciences, 105(44), 17046–17049. https://doi.org/10.1073/pnas.0806560105

30. Hombría, J. C.-G., Brown, S., Häder, S., & Zeidler, M. P. (2005). Characterisation of Upd2, a Drosophila JAK/STAT pathway ligand. Developmental Biology, 288(2), 420–433. https://doi.org/doi.org/10.1016/j.ydbio.2005.09.040

31. Hudson, A. M., & Cooley, L. (2014). Methods for studying oogenesis. Methods (San Diego, Calif)., 68(1), 207–217. https://doi.org/10.1016/j.ymeth.2014.01.005

32. Huypens, P., Sass, S., Wu, M., Dyckhoff, D., Tschöp, M., Theis, F., Marschall, S., de Angelis, M. H., & Beckers, J. (2016). Epigenetic germline inheritance of diet-induced obesity and insulin resistance. Nature Genetics, 48(5), 497–499. https://doi.org/10.1038/ng.3527

33. Ikeya, T., Galic, M., Belawat, P., Nairz, K., & Hafen, E. (2002). Nutrient-Dependent Expression of Insulin-like Peptides from Neuroendocrine Cells in the CNS Contributes to Growth Regulation in Drosophila. Current Biology, 12(15), 1293–1300. https://doi.org/10.1016/S0960-9822(02)01043-6

34. Inagaki, Hidehiko K., Ben-Tabou de-Leon, S., Wong, A. M., Jagadish, S., Ishimoto, H., Barnea, G., Kitamoto, T., Axel, R., & Anderson, David J. (2012). Visualizing Neuromodulation In Vivo: TANGO-Mapping of Dopamine Signaling Reveals Appetite Control of Sugar Sensing. Cell, 148(3), 583–595. https://doi.org/10.1016/j.cell.2011.12.022

35. Ja William, W., Carvalho Gil, B., Mak Elizabeth, M., de la Rosa Noelle, N., Fang Annie, Y., Liong Jonathan, C., Brummel, T., & Benzer, S. (2007). Prandiology of Drosophila and the CAFE assay. Proceedings of the National Academy of Sciences, 104(20), 8253–8256. https://doi.org/10.1073/pnas.0702726104

36. Ja, W. W., Carvalho, G. B., Mak, E. M., de la Rosa, N. N., Fang, A. Y., Liong, J. C., Brummel, T., & Benzer, S. (2007). Prandiology of Drosophila and the CAFE assay. Proceedings of the National Academy of Sciences, 104(20), 8253–8256. https://doi.org/10.1073/pnas.0702726104

37. Karunakar, P., Bhalla, A., & Sharma, A. (2019). Transgenerational inheritance of cold temperature response in Drosophila. FEBS Letters, 593(6), 594–600. https://doi.org/10.1002/1873-3468.13343

38. Kaspar, D., Hastreiter, S., Irmler, M., Hrabé de Angelis, M., & Beckers, J. (2020). Nutrition and its role in epigenetic inheritance of obesity and diabetes across generations. Mammalian Genome, 31(5), 119–133. https://doi.org/10.1007/s00335-020-09839-z

39. Kelly, B. D. (2019). The Great Irish Famine (1845–52) and the Irish asylum system: remembering, forgetting, and remembering again. Irish Journal of Medical Science (1971 -), 188(3), 953–958. https://doi.org/10.1007/s11845-019-01967-z

40. Le Thomas, A., Stuwe, E., Li, S., Du, J., Marinov, G., Rozhkov, N., Chen, Y. C., Luo, Y., Sachidanandam, R., Toth, K. F., Patel, D., & Aravin, A. A. (2014). Transgenerationally inherited piRNAs trigger piRNA biogenesis by changing the chromatin of piRNA clusters and inducing precursor processing. Genes Dev, 28(15), 1667–1680. https://doi.org/10.1101/gad.245514.114

41. Li, C., & Lumey, L. H. (2017). Exposure to the Chinese famine of 1959–61 in early life and long-term health conditions: a systematic review and meta-analysis. International Journal of Epidemiology, 46(4), 1157–1170. https://doi.org/10.1093/ije/dyx013

42. Li, J., Na, L., Ma, H., Zhang, Z., Li, T., Lin, L., Li, Q., Sun, C., & Li, Y. (2015). Multigenerational effects of parental prenatal exposure to famine on adult offspring cognitive function. Scientific Reports, 5(1), 13792. https://doi.org/10.1038/srep13792

43. Li, X. Y., Harrison, M. M., Villalta, J. E., Kaplan, T., & Eisen, M. B. (2014). Establishment of regions of genomic activity during the Drosophila maternal to zygotic transition. Elife, 3. https://doi.org/10.7554/eLife.03737

44. Loh, C. H., van Genesen, S., Perino, M., Bark, M. R., & Veenstra, G. J. C. (2021). Loss of PRC2 subunits primes lineage choice during exit of pluripotency. Nature Communications, 12(1), 6985. https://doi.org/10.1038/s41467-021-27314-4

45. Lyko, F., Ramsahoye, B. H., & Jaenisch, R. (2000). DNA methylation in Drosophila melanogaster. Nature, 408(6812), 538–540. https://doi.org/10.1038/35046205

46. Malik, V. S., & Hu, F. B. (2022). The role of sugar-sweetened beverages in the global epidemics of obesity and chronic diseases. Nature Reviews Endocrinology, 18(4), 205–218. https://doi.org/10.1038/s41574-021-00627-6

47. Malik, V. S., Popkin, B. M., Bray, G. A., Després, J.-P., Willett, W. C., & Hu, F. B. (2010). Sugar-Sweetened Beverages and Risk of Metabolic Syndrome and Type 2 Diabetes: A meta-analysis. Diabetes Care, 33(11), 2477–2483. https://doi.org/10.2337/dc10-1079

48. Margueron, R., & Reinberg, D. (2011). The Polycomb complex PRC2 and its mark in life. Nature, 469(7330), 343–349. https://doi.org/10.1038/nature09784

49. Masuyama, H., & Hiramatsu, Y. (2012). Effects of a High-Fat Diet Exposure in Utero on the Metabolic Syndrome-Like Phenomenon in Mouse Offspring through Epigenetic Changes in Adipocytokine Gene Expression. Endocrinology, 153(6), 2823–2830. https://doi.org/10.1210/en.2011-2161

50. May, C. E., Vaziri, A., Lin, Y. Q., Grushko, O., Khabiri, M., Wang, Q.-P., Holme, K. J., Pletcher, S. D., Freddolino, P. L., Neely, G. G., & Dus, M. (2019). High Dietary Sugar Reshapes Sweet Taste to Promote Feeding Behavior in Drosophila melanogaster. Cell Reports, 27(6), 1675–1685.e1677. https://doi.org/10.1016/j.celrep.2019.04.027

51. Miska, E. A., & Ferguson-Smith, A. C. (2016). Transgenerational inheritance: Models and mechanisms of non–DNA sequence–based inheritance. Science, 354(6308), 59–63. https://doi.org/10.1126/science.aaf4945

52. Mlodzik, M., & Gehring, W. J. (1987). Expression of the caudal gene in the germ line of Drosophila: Formation of an RNA and protein gradient during early embryogenesis. Cell, 48(3), 465–478. https://doi.org/10.1016/0092-8674(87)90197-8

53. Moore, R. S., Kaletsky, R., Lesnik, C., Cota, V., Blackman, E., Parsons, L. R., Gitai, Z., & Murphy, C. T. (2021). The role of the Cer1 transposon in horizontal transfer of transgenerational memory. Cell, 184(18), 4697–4712.e4618. https://doi.org/10.1016/j.cell.2021.07.022

54. Nässel, D., Kubrak, O., Liu, Y., Luo, J., & Lushchak, O. (2013). Factors that regulate insulin producing cells and their output in Drosophila [Review]. Frontiers in Physiology, 4. https://doi.org/10.3389/fphys.2013.00252

55. Nekrasov, M., Klymenko, T., Fraterman, S., Papp, B., Oktaba, K., Köcher, T., Cohen, A., Stunnenberg, H. G., Wilm, M., & Müller, J. (2007). Pcl-PRC2 is needed to generate high levels of H3-K27 trimethylation at Polycomb target genes. The EMBO journal, 26(18), 4078–4088. https://doi.org/10.1038/sj.emboj.7601837

56. Öst, A., Lempradl, A., Casas, E., Weigert, M., Tiko, T., Deniz, M., Pantano, L., Boenisch, U., Itskov, Pavel M., Stoeckius, M., Ruf, M., Rajewsky, N., Reuter, G., Iovino, N., Ribeiro, C., Alenius, M., Heyne, S., Vavouri, T., & Pospisilik, J. A. (2014). Paternal Diet Defines Offspring Chromatin State and Intergenerational Obesity. Cell, 159(6), 1352–1364. https://doi.org/10.1016/j.cell.2014.11.005

57. Painter, R. C., de Rooij, S. R., Bossuyt, P. M., Phillips, D. I., Osmond, C., Barker, D. J., Bleker, O. P., & Roseboom, T. J. (2006). Blood pressure response to psychological stressors in adults after prenatal exposure to the Dutch famine. Journal of Hypertension, 24(9). https://doi.org/10.1097/01.hjh.0000242401.45591.e7

58. Palanker Musselman, L., Fink, J. L., Narzinski, K., Ramachandran, P. V., Sukumar Hathiramani, S., Cagan, R. L., & Baranski, T. J. (2011). A high-sugar diet produces obesity and insulin resistance in wild-type Drosophila. Disease Models & Mechanisms, 4(6), 842–849. https://doi.org/10.1242/dmm.007948

59. Park, J.-H., Kim, S.-H., Lee, M. S., & Kim, M.-S. (2017). Epigenetic modification by dietary factors: Implications in metabolic syndrome. Molecular Aspects of Medicine, 54, 58–70. https://doi.org/10.1016/j.mam.2017.01.008

60. Pendergast, J. S., Branecky, K. L., Yang, W., Ellacott, K. L. J., Niswender, K. D., & Yamazaki, S. (2013). High-fat diet acutely affects circadian organisation and eating behavior. European Journal of Neuroscience, 37(8), 1350–1356. https://doi.org/10.1111/ejn.12133

61. Peng, J. C., Valouev, A., Liu, N., & Lin, H. (2016). Piwi maintains germline stem cells and oogenesis in Drosophila through negative regulation of Polycomb group proteins. Nature Genetics, 48(3), 283–291. https://doi.org/10.1038/ng.3486

62. Pool, A. H., & Scott, K. (2014). Feeding regulation in Drosophila. Curr Opin Neurobiol, 29, 57–63. https://doi.org/10.1016/j.conb.2014.05.008

63. Porte, D., Jr., Baskin, D. G., & Schwartz, M. W. (2002). Leptin and insulin action in the central nervous system. Nutr Rev, 60(10 Pt 2), S20–29; discussion S68-84, 85-27. https://doi.org/10.1301/002966402320634797

64. Qi, W., Wang, G., & Wang, L. (2021). A novel satiety sensor detects circulating glucose and suppresses food consumption via insulin-producing cells in Drosophila. Cell Research, 31(5), 580–588. https://doi.org/10.1038/s41422-020-00449-7

65. Qi, W., Yang, Z., Lin, Z., Park, J.-Y., Suh, G. S. B., & Wang, L. (2015). A quantitative feeding assay in adult Drosophila reveals rapid modulation of food ingestion by its nutritional value. Molecular Brain, 8(1), 87. https://doi.org/10.1186/s13041-015-0179-x

66. Rajan, A., & Perrimon, N. (2012). Cytokine Unpaired 2 Regulates Physiological Homeostasis by Remotely Controlling Insulin Secretion. Cell, 151(1), 123–137. https://doi.org/10.1016/j.cell.2012.08.019

67. Ravelli, A. C. J., van der Meulen, J. H. P., Michels, R. P. J., Osmond, C., Barker, D. J. P., Hales, C. N., & Bleker, O. P. (1998). Glucose tolerance in adults after prenatal exposure to famine. The Lancet, 351(9097), 173–177. https://doi.org/10.1016/S0140-6736(97)07244-9

68. Rechavi, O., Houri-Ze’evi, L., Anava, S., Goh, Wee Siong S., Kerk, Sze Y., Hannon, Gregory J., & Hobert, O. (2014). Starvation-Induced Transgenerational Inheritance of Small RNAs in C. elegans. Cell, 158(2), 277–287. https://doi.org/10.1016/j.cell.2014.06.020

69. Ryu, J.-H., Kim, S.-H., Lee, H.-Y., Bai, J. Y., Nam, Y.-D., Bae, J.-W., Lee, D. G., Shin, S. C., Ha, E.-M., & Lee, W.-J. (2008). Innate Immune Homeostasis by the Homeobox Gene Caudal and Commensal-Gut Mutualism in Drosophila. Science, 319(5864), 777–782. https://doi.org/10.1126/science.1149357

70. Schulz, L. C. (2010). The Dutch Hunger Winter and the developmental origins of health and disease. Proceedings of the National Academy of Sciences, 107(39), 16757–16758. https://doi.org/10.1073/pnas.1012911107

71. Skvortsova, K., Iovino, N., & Bogdanovic, O. (2018). Functions and mechanisms of epigenetic inheritance in animals. Nat Rev Mol Cell Biol, 19(12), 774–790. https://doi.org/10.1038/s41580-018-0074-2

72. Somer, R. A., & Thummel, C. S. (2014). Epigenetic inheritance of metabolic state. Current Opinion in Genetics & Development, 27, 43–47. https://doi.org/10.1016/j.gde.2014.03.008

73. Stegemann, R., & Buchner, D. A. (2015). Transgenerational inheritance of metabolic disease. Seminars in Cell & Developmental Biology, 43, 131–140. https://doi.org/10.1016/j.semcdb.2015.04.007

74. Sung, H., Vesela, I., Driks, H., Ferrario, C. R., Mistretta, C. M., Bradley, R. M., & Dus, M. (2022). High-sucrose diet exposure is associated with selective and reversible alterations in the rat peripheral taste system. Current Biology, 32(19), 4103–4113.e4104. https://doi.org/10.1016/j.cub.2022.07.063

75. Tie, F., Banerjee, R., Stratton, C. A., Prasad-Sinha, J., Stepanik, V., Zlobin, A., Diaz, M. O., Scacheri, P. C., & Harte, P. J. (2009). CBP-mediated acetylation of histone H3 lysine 27 antagonizes Drosophila Polycomb silencing. Development, 136(18), 3131–3141. https://doi.org/10.1242/dev.037127

76. Tobi, E. W., Goeman, J. J., Monajemi, R., Gu, H., Putter, H., Zhang, Y., Slieker, R. C., Stok, A. P., Thijssen, P. E., Müller, F., van Zwet, E. W., Bock, C., Meissner, A., Lumey, L. H., Eline Slagboom, P., & Heijmans, B. T. (2014). DNA methylation signatures link prenatal famine exposure to growth and metabolism. Nature Communications, 5(1), 5592. https://doi.org/10.1038/ncomms6592

77. Tracey, W. D., Jr., Ning, X., Klingler, M., Kramer, S. G., & Gergen, J. P. (2000). Quantitative analysis of gene function in the Drosophila embryo. Genetics, 154(1), 273–284. https://doi.org/10.1093/genetics/154.1.273

78. Vahid, F., Zand, H., Nosrat–Mirshekarlou, E., Najafi, R., & Hekmatdoost, A. (2015). The role dietary of bioactive compounds on the regulation of histone acetylases and deacetylases: A review. Gene, 562(1), 8–15. https://doi.org/10.1016/j.gene.2015.02.045

79. van Dam, E., van Leeuwen, L. A. G., dos Santos, E., James, J., Best, L., Lennicke, C., Vincent, A. J., Marinos, G., Foley, A., Buricova, M., Mokochinski, J. B., Kramer, H. B., Lieb, W., Laudes, M., Franke, A., Kaleta, C., & Cochemé, H. M. (2020). Sugar-Induced Obesity and Insulin Resistance Are Uncoupled from Shortened Survival in Drosophila. Cell Metabolism, 31(4), 710–725.e717. https://doi.org/10.1016/j.cmet.2020.02.016

80. Vaziri, A., Khabiri, M., Genaw, B. T., May, C. E., Freddolino, P. L., & Dus, M. (2020). Persistent epigenetic reprogramming of sweet taste by diet. Science Advances, 6(46), eabc8492. https://doi.org/10.1126/sciadv.abc8492

81. Wan, Q.-L., Meng, X., Wang, C., Dai, W., Luo, Z., Yin, Z., Ju, Z., Fu, X., Yang, J., Ye, Q., Zhang, Z.-H.,& Zhou, Q. (2022). Histone H3K4me3 modification is a transgenerational epigenetic signal for lipid metabolism in Caenorhabditis elegans. Nature Communications, 13(1), 768. https://doi.org/10.1038/s41467-022-28469-4

82. Wang, C., & Lin, H. (2021). Roles of piRNAs in transposon and pseudogene regulation of germline mRNAs and lncRNAs. Genome Biology, 22. https://doi.org/10.1186/s13059-020-02221-x

83. Wang, X., & Moazed, D. (2017). DNA sequence-dependent epigenetic inheritance of gene silencing and histone H3K9 methylation. Science, 356(6333), 88–91. https://doi.org/10.1126/science.aaj2114

84. Wei, Y., Yang, C.-R., Wei, Y.-P., Zhao, Z.-A., Hou, Y., Schatten, H., & Sun, Q.-Y. (2014). Paternally induced transgenerational inheritance of susceptibility to diabetes in mammals. Proceedings of the National Academy of Sciences, 111(5), 1873–1878. https://doi.org/10.1073/pnas.1321195111

85. Wu, Z., Isik, M., Moroz, N., Steinbaugh, M. J., Zhang, P., & Blackwell, T. K. (2019). Dietary Restriction Extends Lifespan through Metabolic Regulation of Innate Immunity. Cell Metabolism, 29(5), 1192–1205.e1198. https://doi.org/10.1016/j.cmet.2019.02.013

86. Xia, B., Gerstin, E., Schones, D. E., Huang, W., & Steven de Belle, J. (2016). Transgenerational programming of longevity through E(z)-mediated histone H3K27 trimethylation in Drosophila. Aging (Albany NY), 8(11), 2988–3008. https://doi.org/10.18632/aging.101107

87. Yang, Z., Huang, R., Fu, X., Wang, G., Qi, W., Mao, D., Shi, Z., Shen, W. L., & Wang, L. (2018). A post-ingestive amino acid sensor promotes food consumption in Drosophila. Cell Research, 28(10), 1013–1025. https://doi.org/10.1038/s41422-018-0084-9

88. Yu, Y., Huang, R., Ye, J., Zhang, V., Wu, C., Cheng, G., Jia, J., & Wang, L. (2016). Regulation of starvation-induced hyperactivity by insulin and glucagon signaling in adult Drosophila. eLife, 5, e15693. https://doi.org/10.7554/eLife.15693

89. Zenk, F., Loeser, E., Schiavo, R., Kilpert, F., Bogdanović, O., & Iovino, N. (2017). Germ line–inherited H3K27me3 restricts enhancer function during maternal-to-zygotic transition. Science, 357(6347), 212–216. https://doi.org/10.1126/science.aam5339

